# A hierarchical GBP network promotes cytosolic LPS recognition and sepsis

**DOI:** 10.1101/2021.08.25.457662

**Authors:** Eui-Soon Park, Bae-Hoon Kim, Pradeep Kumar, Kyle Tretina, Agnieszka Maminska, William M. Philbrick, Richard A. Flavell, John D. MacMicking

**Affiliations:** Howard Hughes Medical Institute, Yale University School of Medicine; New Haven, CT 06510. USA; Yale Systems Biology Institute, West Haven, C.T. 06477. USA; Departments of Immunobiology, Yale University School of Medicine, New Haven, C.T. 06510. U.S.A.; Microbial Pathogenesis, Yale University School of Medicine, New Haven, C.T. 06510. U.S.A.; Department of Internal Medicine, Yale University School of Medicine, New Haven, C.T. 06510. U.S.A.

## Abstract

Bacterial lipopolysaccharide (LPS) is one of the most bioactive substances known. Trace amounts trigger robust immunity to infection but also life-threatening sepsis causing millions of deaths each year. LPS contamination of the cytosol elicits a caspase-dependent inflammasome pathway promoting cytokine release and host cell death. Here, we report an immune GTPase network controls multiple steps in this pathway by genome-engineering mice to lack 7 different guanylate-binding proteins (GBPs). *Gbp2^-^/^-^* and *Gbp3^-^/^-^* mice had severe caspase-11-driven defects that protected them from septic shock. Gbp2 recruited caspase-11 for LPS recognition whereas Gbp3 assembled and trafficked the pyroptotic pore-forming protein, gasdermin D, after caspase-11 cleavage. Together, our results identify a new functional hierarchy wherein different GBPs choreograph sequential steps in the non-canonical inflammasome pathway to control Gram-negative sepsis.

**One-Sentence Summary:** Immune GTPase network orchestrates hierarchical immunity to bacterial products *in vivo*

## Main Text

Metazoans defend against infection by first recognizing microbial structures or pathogen-induced host damage to mount both cell-autonomous and paracrine immune responses (*1, 2*). A powerful trigger of these responses is lipopolysaccharide (LPS), the major constituent of Gram-negative bacterial outer cell walls (*3, 4*). Picomolar amounts of LPS mobilize robust antimicrobial programs, however, systemic exposure causes Gram-negative sepsis with high mortality rates in the human population (*3–7*). Indeed, sepsis accounts for ∼6 million deaths and 30 million cases each year, prompting urgent calls for a better understanding of how LPS triggers this life-threating inflammatory cascade (*7*).

In mammals, LPS is recognized by the host surface TLR4-MD2 receptor complex (*3, 4*) and a subset of cytosolic caspases (CASPASE-4/5 in humans; caspase-11 in mice) (*8–10*). Binding intracellular LPS (iLPS) activates these caspases as part of a “non-canonical” inflammasome pathway that proteolytically cleaves the pore-forming protein, Gasdermin D (GSDMD), to induce pyroptotic cell death and release inflammatory cytokines such as interleukin-1 beta (IL-1β) during endotoxic shock (*11, 12*). How these intracellular events are co-ordinated and the cofactors involved remain important outstanding questions, especially *in vivo*.

We tackled this question using large-scale genome engineering to produce multiple new mouse strains lacking different members of an immune GTPase family termed guanylate-binding proteins (GBPs) (*13*). GBPs were among the first non-core proteins discovered to regulate inflammasome responses to cytosolic bacteria *in vitro* and *in vivo* (*14*). *En bloc* deletion of a 166-kilobase fragment harboring 5 of 11 *Gbp* loci on mouse chromosome 3H1 suggests they also participate in the iLPS recognition pathway as well (*15*). How these new proteins function mechanistically and whether different members perform specialized tasks as part of a novel hierarchical network is currently untested at an organismal level.

We therefore disrupted each *Gbp* gene in the 3H1 cluster via CRISPR-Cas9 editing plus homologous recombineering to produce complete loss-of-function mutations yielding *Gbp1^-^/^-^*, *Gbp2^-^/^-^*, *Gbp3^-^/^-^*, *Gbp5^-^/^-^* and *Gbp7^-^/^-^* mice (Fig. 1A and fig. S1)(*13, 14*). In addition, we extinguished two new adjacent *Gbp* loci on mouse chromosome 5E5 - *Gbp6* and *Gbp10* - that share 99.1% nucleotide identity and are implicated in antibacterial defense (*13*). Each knockout was maintained on a C57BL/6NJ background that otherwise express all 11 Gbps (*13, 16*)(Fig. 1A). It also fixed *Casp11, Dock2* and *Nlrp1b* loci to ensure Gbp-related inflammasome responses were strain- and line-independent (*8,17-18*).

**Figure 1.**
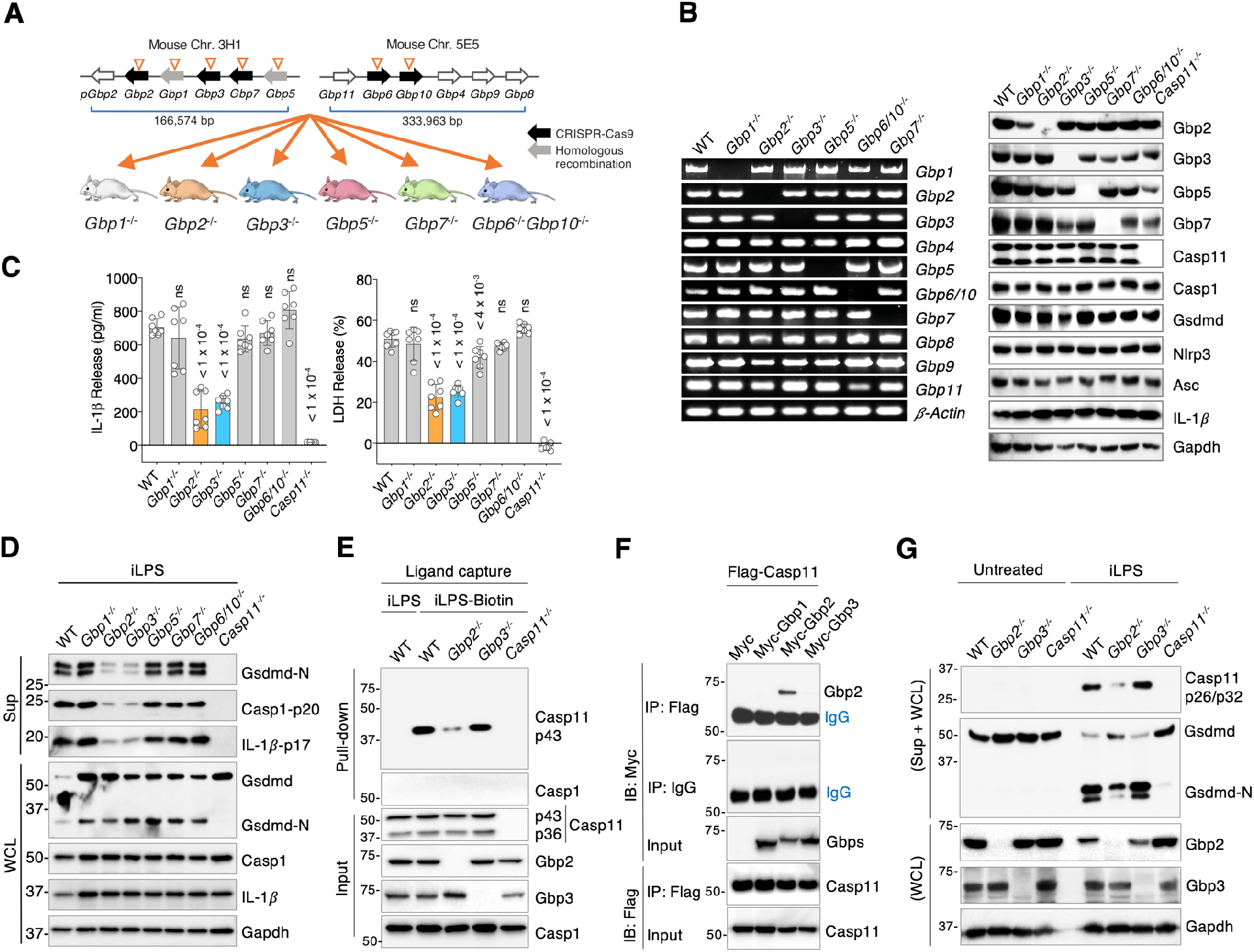
Disruption of 7 *Gbp* loci reveal specific members block the iLPS sensing pathway. **(A)**. *Gbp* knockout mice created by CRISPR-Cas9 editing or homologous recombination. (**B**) Specific loss of *Gbp* mRNA and protein in iLPS-treated BMDMs for each genotype. **(C).** IL-1β and LDH release (mean ± SD) assayed across all Gbp-deficient BMDMs at the same time. *P* values, one-way ANOVA with Dunnett’s multiple comparison test versus wild-type (WT) controls. ns, not significant. Representative of 2 independent experiments across all genotypes and at least 10 independent experiments versus WT controls (fig. S2). **(D).** Gsdmd-N, IL-1β p17 and caspase-1 p20 immunoblotted in supernatants (Sup) and whole cell lysates (WCL) of iLPS-treated BMDMs. Representative of 3 independent repeats. **(E).** iLPS-biotin capture of caspase-11 in Neutravidin pull-down assays. Representative of 3 independent repeats. **(F).** Selective interaction of caspase-11 with Gbp2 but not Gbp1 or Gbp3 in coIP assays. **(G).** Endogenous caspase-11 and Gsdmd cleavage in iLPS-treated BMDMs from different knockout mice. One of 3 similar experiments.

Chromosomal ablation of these 7 *Gbp* loci led to selective loss of *Gbp* mRNA and protein expression using monospecific antibodies where available in LPS-primed bone marrow-derived macrophages (BMDMs). The latter included monospecific antibodies we successfully derived for Gbp3 and Gbp7 in this study plus commercial antibodies for Gbp2 (M15) and Gbp5 (G12) (*13–15*)(Fig. 1B). Importantly, LPS-induced expression of all non-canonical inflammasome components along with the remaining Gbps were intact in these new gene-targeted animals (Fig. 1B).

LPS-primed BMDMs challenged with iLPS as a caspase-11-specific ligand (*5, 6*) revealed pronounced defects in IL-1β release and pyroptosis in cells from *Gbp2^-^/^-^* and *Gbp3^-^/^-^* but not *Gbp1^-^/^-^*, *Gbp5^-^/^-^*, *Gbp7^-^/^-^* and *Gbp6^-^/^-^Gbp10^-^/^-^* mice versus C57BL/6NJ wild-type (WT) controls (Fig. 1C). Defects were evident in multiple experiments spanning BMDMs from age-, sex-, and diet-matched mice at the same time in their diurnal cycle and serologically free of 18 common murine pathogens (Fig. 1B and fig. S2, A and B); hence occult infection, circadian bias, dietary microflora and gender differences were discounted. Moreover, genetic complementation of *Gbp2^-^/^-^* and *Gbp3^-^/^-^* BMDMs with retrovirus expressing Flag-Gbp2 or Flag-HA-Gbp3 rescued IL-1β and LDH release, whereas retrovirus expressing empty plasmid did not (fig. S3A). Thus, phenotypes were attributable to the introduced CRISPR-Cas9 mutations.

Pronounced defects also arose in *Gbp2^-^/^-^* and *Gbp3^-^/^-^* BMDMs when iLPS was cytosolically delivered via electroporation rather than transfection (*5, 10*) or after priming with Pam_3_CSK instead of LPS to induce expression of caspase-11 and GBPs (*8-10,13*) (fig. S3B). Similar outcomes were observed following exposure to *E. coli* outer membrane vesicles (OMVs)(*19*) or *Salmonella enterica typhimurium* SifA mutant (*Stm^ΔSifA^*) that engage the caspase-11-dependent pathway in the cytosol (*20*), but not to isolated agonists of canonical inflammasomes (fig. S3, C to E). Notably, *Gbp2^-^/^-^* and *Gbp3^-^/^-^* macrophages had intact responses to apoptotic stimuli such as H_2_O_2_, necroptotic stimuli like LPS/zVAD, and robust LPS-induced secretion of Tlr4-dependent cytokines such as IL-6 (fig. S3F). Hence Gbp2 and Gbp3 do not broadly regulate cell death or cytokine secretion but were preferentially engaged in inflammasome responses to microbial stimuli. Other priming agents (interferon-gamma; IFN-γ) or bacterial species could conceivably expand this profile to include Gbp1 or Gbp5 (*13–14*). However, Gbp2 and Gbp3 appeared most critical in response to *E. coli* iLPS as a single well-defined caspase-11 agonist.

Following iLPS detection, caspase-11 cleaves gasdermin D to generate ∼30kDa N-terminal subunits (Gsdmd-N) that assemble into 16-28 subunit pores at the plasma membrane (PM) for mature IL-1β export and osmotic rupture during pyroptosis (*11-12,21-23*). Examination of 7 Gbp deficiencies found *Gbp2^-^/^-^* and *Gbp3^-^/^-^* BMDMs had defective Gsdmd-N, mature IL-1β p17 and caspase-1 p20 export in response to iLPS (Fig. 1D), consistent with bioassays measuring IL-1β production and pyroptosis.

Where do these two Gbps act? Early work discovered Gbps target intracellular bacteria shortly after uptake in activated mouse macrophages (*13*). Super-resolution imaging (structured illumination microscopy; SIM) and high-throughput unbiased analysis of 11,804 BMDMs found ∼16.7% of iLPS was targeted by Gbp2 within 15-30 minutes of uptake (fig. S4A). In contrast, Gbp3 co-localization was lower (< 9.3%) in 7,298 BMDMs examined; such targeting was abolished in *Gbp2^-^/^-^* cells, revealing Gbp2 acts upstream of Gbp3 (fig. S4B and E). Notably, Gbp2-coated iLPS lacked the early endosomal Rab5a effector, EEA1 (*19*), indicating Gbp2 converges on iLPS once the ligand enters the cytosol. Here it may recruit caspase-11 for recognition and activation by iLPS.

We tested endogenous caspase-11 binding to iLPS-biotin using Neutravidin capture assays (fig. S5A). Loss of Gbp2 but not Gbp3 impaired caspase-11 binding to iLPS via its CARD-containing p43 domain (Fig. 1E). Gbp2 also bound caspase-11 in cell lysates as well as in cell-free assays (Fig. 1F and fig. S5B). Thus, Gbp2 interacts with and helps recruit caspase-11 to its ligand. Endogenous Gbp2 but not Gbp3 was also captured by iLPS-biotin, albeit weakly since caspase-11 interactions predominate as the *bone fide* LPS sensor (fig. S5A). Notably, loss of caspase-11 binding to iLPS in *Gbp2^-^/^-^* BMDMs was restored by genetic complementation with wild-type Gbp2 but not catalytically inactive Gbp2^K51A/S52A^ that fails to traffic to the ligand (*24*)(Fig. S4D and fig. S5C). Thus, GTPase activity of Gbp2 helps recruit caspase-11 for cytosolic LPS recognition.

Defective ligand binding by caspase-11 in *Gbp2^-^/^-^* BMDMs resulted in loss of caspase-11 autoactivation (denoted by the p26 fragment) and subsequent cleavage of its Gsdmd substrate to yield Gsdmd-N (Figs. 1G and 2A, fig S5D and E). Nlrp3-Asc foci formation was consequently impaired in 15,313 *Gbp2^-^/^-^* BMDMs examined because Gsdmd-N pores allow potassium efflux for Nlrp3 inflammasome activation (*11, 12*)(Fig.2A, fig S5F and G). Despite normal caspase-11 activity, defects in Nlrp3-Asc foci formation were also evident across 19,014 *Gbp3^-^/^-^* BMDMs (fig S5F and G), consistent with earlier IL-1β and pyroptotic defects. This positioned Gbp3 downstream of iLPS recognition but upstream of Nlrp3 inflammasome assembly (Fig. 2A). Gbp2 and Gbp3 could therefore act sequentially as part of a new hierarchical network.

**Figure 2.**
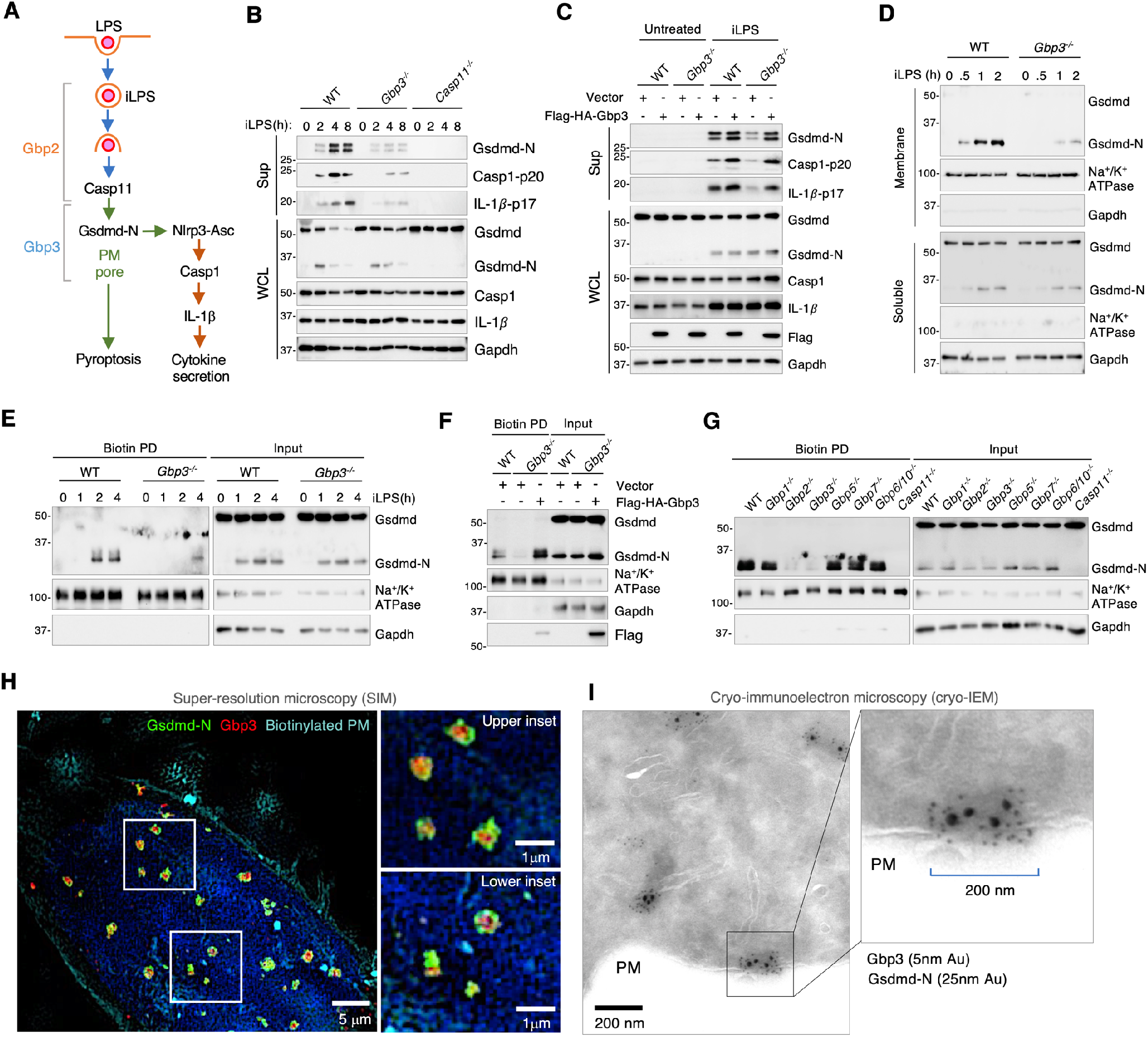
Lost PM targeting by Gsdmd-N in *Gbp3^-^/^-^* macrophages. **(A)**. Potential positions of Gbp2 and Gbp3 in the non-canonical inflammasome pathway. **(B)**. Secreted Gsdmd-N, IL-1β p17 and caspase-1 p20 after iLPS exposure in BMDMs. Sup, supernatants. WCL, whole cell lysates. Representative of 3 independent repeats. **(C)**. Gsdmd-N, IL-1β p17 and caspase-1 p20 in genetically complemented *Gbp3^-^/^-^* BMDMs. Representative of 2 independent repeats. **(D)**. Gsdmd-N in BMDM membrane extracts after iLPS exposure. Na^+^/K^+^ ATPase, membrane marker; Gapdh, soluble marker. Representative of 4 independent repeats. **(E)**. Neutravidin pull-down (PD) of biotinylated PM immunoblotted for Gsdmd-N. Input, total cell lysates. Representative of 4 independent repeats. **(F)**. Gsdmd-N in biotinylated PM fractions 2h after iLPS transfection of complemented BMDMs. Input, total cell lysates. Representative of 2 independent repeats. **(G)**. PM-captured Gsdmd-N 2h after iLPS transfection in BMDMs from all 7 Gbp genotypes plus WT and *Casp11^-^/^-^* controls. Representative of 2 independent repeats. **(H)**. SIM imaging of doxycycline-expressed Gsdmd-N and Gbp3 in HeLa cells with PM biotinylation. Insets shown on right. **(I)**. Cryo-immunoelectron microscopy (cryo-IEM) of Gsdmd-N and Gbp3. Sections labelled with anti-RFP and anti-GFP antibodies conjugated to 5nm and 25nm gold (Au) particles, respectively. Inset, right.

We focused on events immediately after caspase-11 activation in an attempt to delineate how Gbp3 operates. Gsdmd-N liberated by caspase-11 cleavage form pores in the PM to release IL-1β p17 along with the active caspase-1 p20 subunit (Fig. 2A). We detected IL-1β p17, caspase-1 p20 and Gsdmd-N in WT supernatants between 2-8 hrs after iLPS treatment; they were largely absent, however, from *Gbp3^-^/^-^* BMDMs, despite intact cleavage of 55kDa Gsdmd to ∼30kDa Gsdmd-N in whole cell lysates (WCL)(Fig. 2B). Such defects were rescued in *Gbp3^-^/^-^* BMDMs by Flag-HA-Gbp3, supporting a role for Gbp3 in controlling this important step (Fig. 2C).

IL-1β export requires that Gsdmd-N assemble on lipid membranes to generate multi-subunit pores (*11-12,21-23*). Gsdmd-N partitioned into membrane extracts within 30-120 min of iLPS exposure that was again greatly diminished in *Gbp3^-^/^-^* BMDMs (Fig. 2D). To test if this loss occurred at the PM, surface biotinylation and cross-linking was used to label endogenous Gsdmd-N for capture by Neutravidin beads. This method successfully retrieved PM-localized Gsdmd-N which was barely detectable in *Gbp3^-^/^-^* BMDMs (Fig. 2E). Genetic complementation in *Gbp3^-^/^-^* cells restored surface Gsdmd-N with Flag-HA-Gbp3 also being present in PM fractions (Fig. 2F). Surface biotinylation assays across 7 Gbp deficiencies reinforced these phenotypes: only *Gbp2^-^/^-^* and *Gbp3^-^/^-^* BMDMs exhibited major loss of PM-localized Gsdmd-N in response to iLPS (Fig. 2G). For Gbp2, this outcome resulted from its upstream involvement in caspase-11 recruitment and activation by iLPS, whereas Gbp3 promotes Gsdmd-N assembly and/or trafficking after cleavage by caspase-11 (Fig. 2A).

To observe where Gbp3 promotes this activity, we developed a doxycyline-regulated bipartite P2A reporter system to avoid constitutive Gsdmd-N cytotoxicity during super resolution imaging (fig. S6A to F). LDH release assays confirmed reporter functionality after doxycycline treatment with co-expression of Gbp3 enhancing Gsdmd-N-dependent pyroptosis (fig. S6B) Strikingly, > 50% of Gbp3 assembled into large “ring-like” structures together with Gsdmd-N inside mammalian cells (Fig. 2H and fig. S7A to C). Gbp2 was essentially absent from these complexes (fig. S7C). Cryo-immunoelectron microscopy (cryo-IEM) found Gbp3-Gsdmd-N structures were ∼100-200nm in diameter (Fig. 2I), large enough to accommodate 30nm Gsdmd-N protomers with 140-180 Å pore interiors (*21, 23*). Many of these cryo-IEM structures were cytosolic. Thus, Gbp3 may facilitate Gsdmd-N pre-assembly before transport to the cell surface (Fig. 2E to G). Such a model fits total membrane extracts at 30-60 min where Gsdmd-N is still predominantly intracellular while surface biotinylation detects this complex at later times, between 2-4 hrs after iLPS exposure (Fig. 2D and E).

Because Gbp3 is a new uncharacterized member of the GBP family, we generated recombinant Gbp3 plus putative mutants to identify biochemical activities needed for its inflammasome-related functions (Fig. 3A, fig. S8A and B). Radioactive [^32^P]-GTP hydrolysis assays revealed murine Gbp3 produces both [^32^P]-GDP and [^32^P]-GMP (Fig. 3A and fig. S8C). In contrast, putative catalytic site (K45A/S46A; S46N) and assembly (D176N) mutants failed to generate [^32^P]-GMP while producing small amounts of [^32^P]-GDP (Fig. 3A and fig. S8C). An R3A C-terminal mutant (Gbp3^R3A^) similar to a human GBP1 motif needed for trafficking (*25*) had intact enzyme activity.

**Figure 3.**
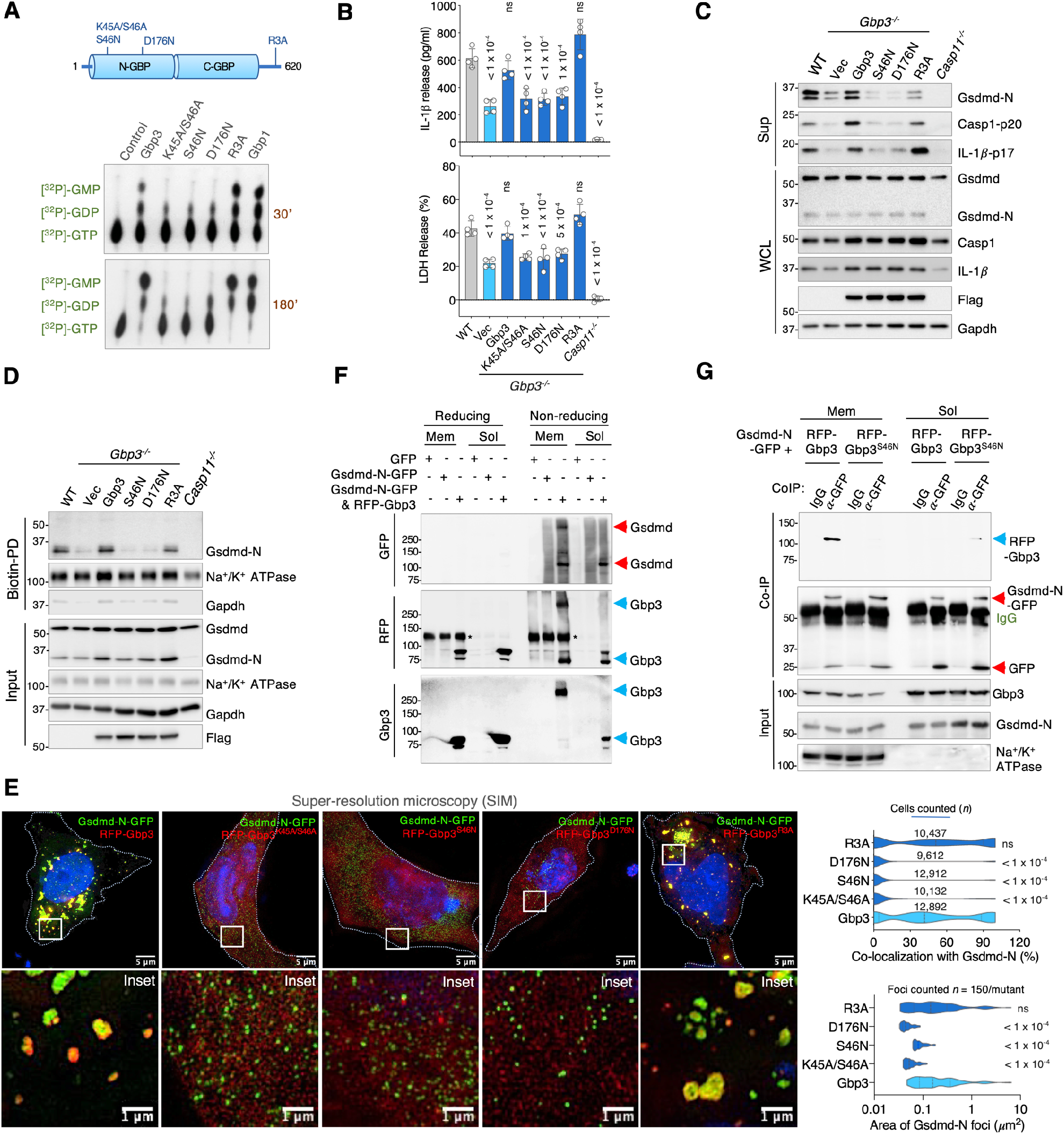
Gbp3 promotes Gsdmd-N assembly *en route* to the PM. **(A)**. Catalytic activity of recombinant Gbp3 and its mutants in radiolabeled GTPase assays at 30 and 180 min after substrate addition. Mutations shown above. **(B)**. IL-1β and LDH release in iLPS-treated *Gbp3^-^/^-^* BMDMs complemented with Gbp3 or its mutants. Mean ± SD. *P* values, one-way ANOVA with Dunnett’s multiple comparison test versus wild-type controls. ns, not significant. **(C)**. Gsdmd-N, IL-1β p17 and caspase-1 p20 in *Gbp3^-^/^-^* BMDMs complemented with Gbp3 or its mutants. Representative of 3 independent repeats. **(D)**. PM-captured Gsdmd-N 2h after iLPS transfection in *Gbp3^-^/^-^* BMDMs complemented with Gbp3 or its mutants. Representative of 3 independent repeats. **(E)**. Gsdmd-N pre-assembly structures in HeLa cells co-expressing Gbp3 or its mutants. Gbp3-Gsdmd-N colocalization and size shown in violin plots (right). *P* values, one-way ANOVA with Dunnett’s multiple comparison test of mutants versus active Gbp3. ns, not significant. **(F)**. Higher molecular weight Gsdmd-N and Gbp3 species detected in membrane and soluble fractions of 293T cells by non-reducing PAGE. Representative of 2 independent repeats. **(G)**. Co-IP of active or inactive Gbp3 by Gsdmd-N in membrane and soluble fractions of 293T cells. IgG, heavy and light immunoglobulin chains. Representative of 3 independent repeats.

Subsequent complementation of iLPS-treated *Gbp3^-^/^-^* BMDMs found mutants that failed to generate GMP (Flag-HA Gbp3^K45A/S46A^, Flag-HA Gbp3^S46N^, Flag-HA Gbp3^D176N^) also failed to rescue IL-1β secretion and pyroptosis, whereas catalytically active Flag-HA Gbp3 or Flag-HA Gbp3^R3A^ both reversed these defects (Fig. 3B). Identical outcomes were observed for Gsdmd-N, caspase-1 p20 and IL-1β p17 release (Fig. 3C) as well as Neutravidin capture of PM Gsdmd-N following surface biotinylation (Fig. 3D). Thus, Gbp3 catalysis and GMP production is essential for its non-canonical inflammasome-related activity.

How does Gbp3 catalysis confer these effects? Super-resolution imaging found catalytically active Gbp3 and Gbp3^R3A^ induced large Gsdmd-N structures *in situ* whereas inactive RFP-Gbp3^K45A/S46N^, RFP-Gbp3^S46N^ and RFP-Gbp3^D176N^ did not (Fig. 3E and fig. S6E). The Stokes radius (R_s_) of catalytically active Gbp3 is ∼10 kDa (98-99 kDa; R_s_, 4.14nm) larger than all of its inactive mutants (87-92 kDa, R_s_, 3.96-4.03nm) (fig. S8D and E). This closely resembles catalytically active human GBP1 that forms elongated conformers (*26*). Adopting an extended conformation could enable Gbp3 to provide a nucleating platform for Gsdmd-N oligomerization and assembly.

We tested this idea by expressing each protein singly or in combination before electrophoretically separating Gsdmd-N complexes *in situ* under non-reducing conditions (*22, 23*). Large >250 kDa Gsdmd-N complexes accumulated in cell membrane extracts when Gbp3 was co-expressed; RFP-Gbp3 co-migrated with Gsdmd-N as detected by anti-Gbp3 or anti-RFP antibodies (Fig. 3F). In contrast, cells expressing inactive RFP-Gbp3^S46N^ failed to promote Gsdmd-N oligomerization or Gbp3 assembly (fig. S9A). Thus, conformational changes in Gbp3 brought about by GTP hydrolysis facilitates Gsdmd-N incorporation into stable macromolecular species. Such Gsdmd-N complexes were also seen after iLPS exposure in WT but not *Gbp3^-^/^-^* BMDM membrane extracts via native PAGE, indicating Gbp3-driven Gsdmd complexes assemble under endogenous conditions as well (fig. S9B). Here the apparent molecular mass (between 488-1,048 kDa) in iLPS-treated BMDMs fits gasdermin pore estimations (∼496-868 kDa) from recent cryoEM studies (*21, 23*).

Whether these complexes arise via direct or indirect Gbp3-gasdermin interactions was next tested via coIP and pulldown assays. Gsdmd-N-GFP co-immunoprecipitated wild-type RFP-Gbp3 but not RFP-Gbp3^S46N^ and only from iLPS-treated membrane but not soluble fractions, fitting the co-migration profile above (Fig. 3G and fig. S9A). Likewise, Gbp3 co-immunoprecipitated a less toxic L193D Gsdmd-N variant that enabled sufficient membrane fractions to be collected for interaction analysis in the reverse direction (fig. S9C). No binding to the Gsdmd C-terminal region and very weak binding to the auto-inhibited full-length Gsdmd protein was observed (fig. S9C). Notably, Ni-NTA pulldown assays and *in vitro* immunoprecipitation assays did not detect direct interactions (fig. S9D); thus, additional iLPS-induced cofactors probably work with Gbp3 to stabilize the membrane-associated Gsdmd-N structure. This may also explain why Gbp3 preferentially impacts the non-canonical versus canonical pathway since iLPS elicits Gbp3 partners to initiate this process. Here Gbp3 works together with Gbp2 to assemble directly onto iLPS to activate caspase-11-dependent cleavage of Gsdmd but does not assemble on canonical inflammasome ligands like dsDNA or flagellin in the cytosol to activate caspase-1 cleavage of the same substrate (data not shown). Hence other host proteins must regulate Gsdmd assembly under those conditions. Importantly, Gbp3 is the first protein required for Gsdmd pre-assembly and export of the pore-forming complex to the cell surface in response to Gram-negative LPS detected within the host cytosol.

Emergence of this unexpected role arose from studying new *Gbp3^-^/^-^* mice. We therefore asked if such mice were protected from caspase-11-dependent sepsis that requires Gsdmd (*5-6,8*). Large-scale challenge of 125 mice across 8 genotypes with polyI:polyC plus LPS found robust survival in *Gbp2^-^/^-^*, *Gbp3^-^/^-^* and *Casp11^-^/^-^* hosts (Fig. 4A). This protection was also evident in measurements of body surface temperature, serum IL-1β levels and Gsdmd-N membrane targeting, indicating the Gsdmd defects observed *in vitro* also operate *in vivo* (Fig. 4B to D, fig. S10A and B). Notably, *Gbp2^-^/^-^*, *Gbp3^-^/^-^* and *Casp11^-^/^-^* mice were still susceptible to Tlr4-dependent sepsis (*5, 6*), underscoring specificity of the caspase-11 response (fig. S10C).

**Figure 4.**
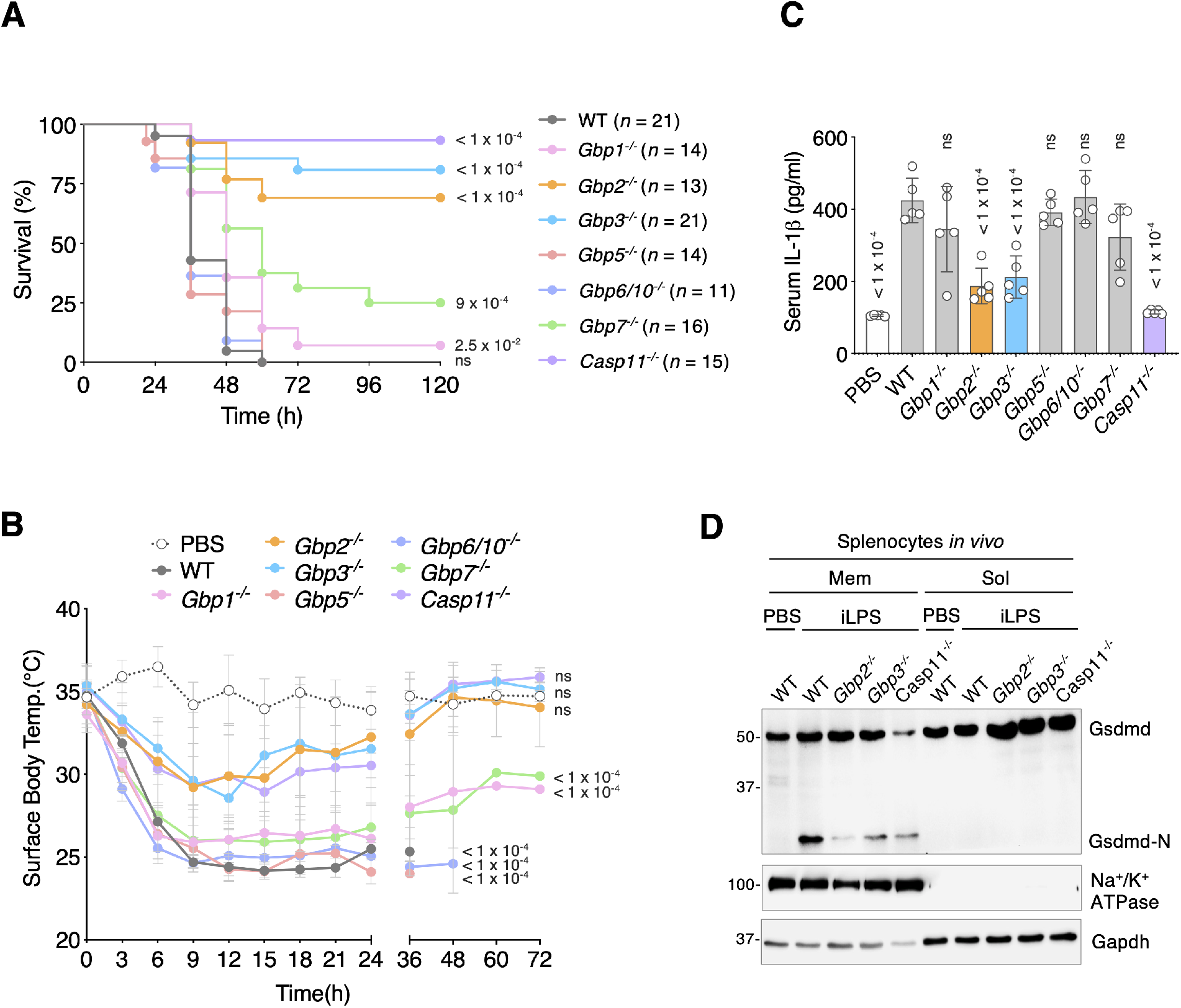
GBP deficiency inhibits Gsdmd-N membrane targeting and sepsis *in vivo*. **(A)**. Kaplan-Meier survival plot of knockout mice to caspase-11-dependent sepsis. *P* values, each knockout group shown versus WT mice using the Gehan-Breslow-Wilcoxon test. *n*, number of mice per group. **(B)**. Surface body temperature monitored every 3 hr up to 24 hr, and each 12 hr thereafter to 72 hr during sepsis. *P* values, for each group using one-way ANOVA with Dunnett’s multiple comparison test versus PBS control. ns, not significant at *P* > 0.05. *n*, 5 mice/group. **(C)** Serum IL-1β at 18 hr p.i. in each sepsis group plus PBS control. *P* values for each knockout group using one-way ANOVA with Dunnett’s multiple comparison test versus wild-type C57BL/6NJ controls. ns, not significant at *P* > 0.05. *n*, 5 mice/group. **(D).** Membrane versus soluble partitioning of endogenous Gsdmd-N in splenocytes at 16 hr after LPS injection. Representative of 2 independent repeats.

Together, our results uncover a new functional hierarchy in the non-canonical inflammasome pathway *in vivo*. Gbp2 promotes ligand detection by caspase-11 while Gbp3 aids Gsdmd-N pre-assembly shortly after caspase-11 cleavage (fig. S10D). Direct LPS coating by mouse Gbp2 and Gbp3 resembles human GBPs 1-4 recently shown to encapsulate the Gram-negative bacterial surface within infected cells (*25,27-28*). Here mouse Gbp2 functions like its closest human ortholog, GBP1, in directly engaging LPS as the proximal Gbp upstream. Whether human GBPs also act analogously to mouse Gbp3 for GSDMD transport, however, is unknown. The present study therefore raises the possibility of new GSDMD trafficking and assembly roles for human GBPs during inflammasome activation. Most importantly, these functions operated within an intact host and suggest that therapeutic inhibition of certain GBPs could potentially help limit tissue damage while preserving the cell-autonomous defense functions of other family members during Gram-negative infection (*29–30*). Our large-scale genome engineering approach thus yields a novel conceptual framework with implications for sepsis and infection control within the human population (*7, 30*).

## Acknowledgements

We thank Dijin Xu and Alex Tunaru for experimental advice and technical help. We are grateful to the Yale Genome Editing Center for gRNA design and microinjection.

## Author contributions

E-S.P. and J.D.M. conceptualized the project and wrote the paper. E-S.P., B-H.K., P.K., K.T., and A.M. performed all experiments with external collaboration by W.P. and R.A.F. to help generate *Gbp2^-^/^-^*, *Gbp3^-^/^-^*, *Gbp7^-^/^-^* and *Gbp6^-^/^-^Gbp10^-^/^-^* mice.

## Data and materials availability

All data needed to evaluate the conclusions are present in the paper or the supplementary materials.

## Supplementary Materials

### MATERIALS & METHODS

#### Cloning and constructs

Murine *Gbp3* (NM_018734) was amplified with indicated tags by PCR from bone marrow-derived macrophage (BMDM) cDNA, and Gbp3 point mutants (K45A/S46A, S46N, D176N and R3A) were generated by using QuikChange site-directed mutagenesis kit (Stratagene) with the mutated oligonucleotides (K45A/S46A: 5’-GTT GGT TTA TAT CGT ACA GGG GCA GCG TAC CTC ATG AAT CGT CTT GCA GGA CGG-3’, S46N: 5’-GTT GGT TTA TAT CGT ACA GGG AAG AAC TAC CTC ATG AAT CGT CTT GCA GGA CGG-3’, D176N: 5’-CCA GAC TTT ATC TGG GCT GTT CGA AAT TTT GCT CTG GAG CTG AAG TTA AAT GGT CGG-3’, R3A: 5’-CAC CTG AGA GAA GAG ATG GAA GCA ACA GCA GCG AAA CCC TCA CTG TTT GG-3’). Murine Gsdmd full-length (Gsdmd), N-terminal fragment (1-248 amino acids), and C-terminal fragment (277-487 amino acids) were amplified with indicated tags by PCR from murine *Gsdmd* (NM_026960) ORF Clone (Origene). Amplicons were cloned into the pMX-IP, pEGFP-n1, pERFP-c1, pCMV-Myc and pET28a plasmids. Gsdmd-N (I105N) and Gsdmd-N (L193D) point mutants were generated by site-directed mutagenesis as mentioned above with the mutated oligonucleotides (I105N: 5’-GCA TGG GAG AAG GGA AAA ATT CTG GTG GGG CTG CAG TGT C-3’, L193D: 5’-CTG CCT GGA GCT TTA TGC GAC AAG GGT GAA GGC AAG GGC C-3’). Flag-Gbp1, Flag-Gbp2, Flag-Gbp5, and Flag-Gbp7 were amplified from BMDM cDNA, and cloned into pMX-IP plasmids. pERFP-c1-Gbp2 and -Gbp5 were cloned from BMDM cDNA. pERFP-c1 plasmids containing Gbp1, or Gbp7, and pMX-IP plasmids containing YFP-Gbp2, or YFP-Gbp2^K51A/A52A^, were as described (*13, 14*). For the doxycycline-inducible system, Gsdmd-N-GFP, RFP-Gbp2, RFP-Gbp3, and RFP-Gbp3 mutants were amplified by PCR from pEGFP-n1 Gsdmd-N or pERFP-c1-Gbp2 or -Gbp3 or -Gbp3 mutants, respectively. Amplicons were cloned into pCW57-MCS1-2A-MCS2 (Addgene). All resultant constructs were sequenced for validation. All the plasmids used in this study are listed in Table S1.

#### Genetically engineered mice

*Gbp2^-/-^*, *Gbp3^-/-^*, *Gbp7^-/-^* and *Gbp6/10^-/-^* mice were generated by CRISPR-Cas9 genome engineering on a C57BL/6NJ background using the gRNA sequences in Table S2 and according to the methods described in Nowarski *et al.*, (*31*). *Gbp6* and *Gbp10* share 99.1% nucleotide identity and were doubly extinguished to avoid functional redundancy. CRISPR-Cas9 ablation of individual or paired *Gbp* loci screened by PCR (fig. S1) were verified by genomic sequencing (Table S3). In addition, loss of mRNA and protein expression was corroborated by qRT-PCR and immunoblotting from bone marrow-derived macrophages (BMDMs). These were taken from progeny mice that received germline CRISPR-Cas9 mutations transmitted from their C57BL/6N knockout parents.

*Gbp1^-/-^* and *Gbp5^-/-^* mice were generated as reported (***13,14***) and backcrossed onto C57BL/6NJ at least 12 generations for this study (> N11). Unlike C57BL/6J mice, both C57BL/6NJ and 129/SvEv strains express all 11 Gbp family members in response to multiple agonists (*13,16,32*). Hence knockout mice are matched across the chromosome 3H1 *Gbp* cluster for functional Gbp expression (Fig. 1), irrespective of whether mutations were introduced by CRISPR-Cas9 editing or homologous recombination.

Likewise, *Casp11^-/-^* mice (*Casp4^tm1Yuan^*; Jackson Laboratory, Bar harbor, MN) on a commercially mixed C57BL/6J and C57BL6/NJ background were also bred with C57BL/6NJ for at least 8 generations. This congenic *Casp11^-/-^*^B6NJ^ subline expresses all 11 Gbps and remaining inflammasome components like parental C57BL/6NJ controls (Fig. 1). In turn, *Gbp1^-/-^*, *Gbp2^-/-^*, *Gbp3^-/-^*, *Gbp5^-/-^*, *Gbp7^-/-^* and *Gbp6^-/-^Gbp10^-/-^* C57BL/6NJ mice express caspase-11 and other inflammasome components like parental C57BL/6NJ mice (Fig. 1). This bidirectional validation along with genetic complementation ensured phenotypes were attributable to the introduced mutations. We purchased *Tlr4^-/-^* (B6.B10ScN-*Tlr4^lps-del^*/JthJ) mice (stock no. 007227) from Jackson lab. All mice used for the derivation of primary BMDMs *and in vivo* experiments were approved by the Yale Institutional Animal Care Committee. They were housed in the same room of a specific-pathogen free (SPF) facility at Yale West Campus.

#### Cell culture

HeLa CCL2, 293T, 293E and L-929 cells (ATCC) were maintained in Dulbecco’s modified Eagle’s medium (DMEM, Gibco) supplemented with 10% fetal bovine serum (FBS, Invitrogen) and 1x Penicillin-Streptomycin solution (Thermo Fisher Scientific) in a 5% CO_2_ at 37°C. BMDMs were isolated from C57BL6/NJ *wild-type*, *Gbp1^-/-^, Gbp2^-/-^, Gbp3^-/-^, Gbp5^-/-^, Gbp6/10^-/-^, Gbp7^-/-^* and *Caspase-11^-/-^* mice. In brief, bone marrow precursor cells were collected from the tibias and femurs of 8-12 weeks old mice. The bone marrow cells were cultured in DMEM supplemented with 10% FBS, and 30% L-929 supernatants in a 5% CO_2_ at 37°C for 5-6 days to differentiate into BMDMs.

#### Semi-quantitative RT-PCR

BMDMs (5×10^6^ cells) were incubated in 10-cm dish with 0.5μg of ultrapure LPS (*E. coli* O111:B4, Invivogen) for 16h. Cells were harvested, and total RNA isolated by RNeasy Mini kit (Qiagen) before quantification by measurement of absorbance at 260nm. Reverse transcription was performed with 2μg of total RNA and M-MLV reverse transcriptase (Thermo Fisher Scientific) for 1h at 42°C followed by 10min at 95°C, and the resulting cDNAs were subjected to semi-quantitative RT-PCR analysis (9800 Fast Thermal Cycler, Applied Biosystems) with indicated primers for *Gbp1 (NM_010259.2), Gbp2 (NM_010260.1), Gbp3 (NM_001289492), Gbp4 (NM_008620.3), Gbp5 (NM_153564.2), Gbp 6 (NM_194336), Gbp7 (NM_145545.3), Gbp8 (NM_029509.4), Gbp9 (NM_172777.4), Gbp10 (NM_001039646.2), Gbp11 (NM_001039647.2)* and *β-actin* (Table S4).

#### Inflammasome activation and treatments

BMDMs were seeded at 5×10^4^ cells/well into 96-well plates or 3×10^5^ cells/well into 24-well plates. For non-canonical inflammasome activation, cells were primed with 0.5μg/ml of ultrapure LPS (*E. coli* O111:B4, Invivogen) or 0.5μg/ml of Pam_3_CSK4 (Invivogen) for 16h, and then transfected with 2μg/ml of ultrapure LPS with 0.5% Fugene HD (Promega) according to manufacturer’s instructions, or treated with 20μg of outer membrane vesicle (OMV) from *E. coli*. At the indicated times, cell supernatants and extracts were collected. For non-canonical inflammasome activation by bacterial infection, *Salmonella typhimurium ΔsifA* (12023, a gift from Dr. D. Holden, Imperial College, London) was grown in LB broth at 37°C overnight. For induction of SPI2 expression, bacteria were incubated with SPI2 induction medium (0.1% w/v casamino acids, 38mM glycerol, 5mM KCl, 0.2mM MgCl_2_, 7.5mM(NH_4_)_2_SO_4_, 0.5mM K_2_SO_4_, 1mM KH_2_PO_4_, 100mM BisTris, 100mM Tris, 100mM Hepes; pH. 6.5) at 37°C for 16-18h. Bacteria were added to BMDMs at MOI 50, centrifuged for 5min at 350x*g*, and then incubated at 37°C for 1h. Extracellular bacteria were killed with 100μg/mL of Gentamycin (Thermo Fisher Scientific) for 1h. After 6h pi, supernatants and cell extracts were collected.

For non-canonical inflammasome activation by LPS electroporation, 3×10^6^ cells of LPS- or PAM3CSK4-primed BMDM were prepared with 5μg of LPS in 2mm gap electroporation cuvette, and electroporated according to manufacturer’s instruction (Protocol #0934, ECM 830 electroporator, BTX). Electroporated BMDMs were collected, and incubated for 1-4h in 37°C CO_2_ incubator. Supernatants and cell lysates were harvested for further analysis.

For canonical inflammasome activation, cells were primed with 0.5μg/ml of ultrapure LPS for 5h, and then treated with either 5mM of ATP (Invivogen) or 250μg/ml of MSU (Invivogen) for 2h. For double-stranded DNA transfection, 2μg of poly(dA:dT) (Invivogen) was transfected using Lipofectamine 2000 (Invitrogen) for 6h according to manufacturer’s instructions. For bacterial flagellin transfection, 70ng of Fla-ST (Standard flagellin from *S. typhimurium*, Invivogen) was transfected using Lipofectamine 2000 for 6h according to manufacturer’s instructions. At the indicated times, supernatants and cell extracts were harvested.

Cell death and IL-1β release was measured by CytoTox 96® Non-Radioactive Cytotoxicity Assay (Promega) and IL-1β ELISA assay kit (Thermo Fisher) according to manufacturer’s instructions, respectively. For NF-κB-mediated inflammatory response, cells were incubated with 0.5μg/ml of ultrapure LPS for 16h, IL-6 level was measured by IL-6 ELISA (BD Bioscience) according to manufacturer’s instructions. To measure necroptotic cell death and apoptotic cell death, cells were incubated with ultrapure 0.1μg of LPS (Invivogen) and 10μM of z-VAD-FMK (R&D systems) with or without 20μM of Necrostatin-1 (Nec-1, Sigma-Aldrich) for 24h, or incubated with 1mM of H_2_O_2_ (Sigma-Aldrich) for 24h, respectively. The levels of cell death were measured as stated above. All assays were performed on a SpectraMax i3x Multimode Plate Reader (Molecular Devices).

#### Retrovirus production and transduction

For retrovirus production, 293T cells were seeded at 5×10^6^ cells/dish in 10-cm dish. pMX-IP vectors containing the gene of interest, pGag/Pol (Addgene), and pCMV-VSV-G (Addgene) were transfected with TransIT-LT1 transfection reagent (Mirus Bio) at 3:1:0.2 ratio. After 48h, supernatants containing retrovirus (retroviral soup) were collected and filtered by 0.45μm PVDF membrane (Millipore). For retroviral transduction, BMDMs were seeded at 5×10^6^ cells/dish in 10-cm petri dish with culture medium (DMEM containing 10% FBS and 30% L-soup). After 24h, cells were incubated with retroviral soup containing 8μg/ml of polybrene (Sigma-Aldrich) for 6-8h, and then retroviral soup was replaced with fresh culture medium. After 16h, BMDMs were detached and resuspended with fresh culture media with 2μg/ml of puromycin (Sigma-Aldrich) to select retrovirus-transduced BMDMs for 2-3days.

#### Intracellular LPS targeting, Asc foci and image analysis

BMDMs were seeded at 5×10^4^ cells/well into 96-well black polystyrene microplate (Sigma-Aldrich), and primed with LPS for 16h. To visualize intracellular LPS, 4μg/ml of LPS-Alexa Fluor™ 594 conjugate (Lipopolysaccharides from *Escherichia coli* Serotype (055:B50), Thermo Fisher Scientific) were transfected. At the indicated times, cells were fixed with 4% paraformaldehyde (SantaCruz Biotechnology) for 15min, permeabilized with 0.2% Triton X-100 for 3min, blocked with 5% normal donkey serum (Sigma-Aldrich) in PBS for 1h, and then incubated with indicated antibodies at 4°C overnight. Cells were incubated with indicated secondary antibodies at RT for 1h, and the nucleus stained by Hoechst 33342 (ThermoFisher Scientific). For intracellular LPS-induced Asc foci analysis, LPS-primed BMDMs were transfected with ultrapure LPS. After indicated times, cells were fixed, permeabilized, and incubated with anti-ASC antibody (D2W8U, Cell Signaling) at 4°C overnight. Thereafter cells were incubated with anti-Rabbit secondary antibody (Rabbit IgG(H+L)-Alexa Fluor 488 conjugate, Thermo Fisher) at RT for 1h, and the nucleus stained by Hoechst 33342. Images were captured on a Nikon Ti Eclipse microscope and z-stacks deconvolved using NIS-Elements Advanced Research software (Nikon); a DeltaVision™ OMX SR microscopy system with structured illumination microscopy (SIM) (GE Healthcare); or IN Cell microscopy (IN Cell Analyzer 2200 Imaging System, GE Healthcare), and analyzed by Image J/Fiji software (Image J 2.0.0). Quantification was analyzed by CellProfiler cell image analysis software (CellProfiler 3.0.0).

#### Co-localization analysis by super resolution microscopy and cryo-IEM

HeLa cells were seeded at 1×10^5^ cells into 24-well plate with 12mm-coverslip (Precision Cover Glasses, #1.5H Thickness, Ø12 mm, ThorLabs), and transfected with doxycycline-inducible pCW57 plasmids containing Gsdmd-N-GFP, Gsdmd-N-GFP/RFP-Gbp2, Gsdmd-N-GFP/RFP-Gbp3 or Gsdmd-N-GFP/RFP-Gbp3 mutants (K45A/S46A, S46N, D176N, and R3A) respectively, by TransIT-LT1 transfection reagent (Mirus Bio) according to manufacturer’s instruction. Cells were treated with 1μg of doxycycline for 24h, fixed, and nuclei were stained. To label plasma membrane, cells were incubated with 2mg/ml of Biotin (EZ-Link™ Sulfo-NHS-LC-LC-Biotin, Thermo Fisher Scientific) at 4°C for 30min, washed with 100mM glycine and 20mM glycine, and incubated with Streptavidin/Alexa Fluor® 647 conjugate (Thermo Fisher) before 4% paraformaldehyde fixation. Super-resolution images were captured on a DeltaVision™ OMX SR microscopy system with structured illumination microcopy (SIM) technique (GE Healthcare), and analyzed by Image J/Fiji software (Image J 2.0.0). For cryo-immunoelectron microscopy (cryo-IEM), HeLa cells were transfected as above and subjected to fixation/rehydration steps and immunogold labeling as described (***13***).

#### Biotinylated-LPS pull-down assay

LPS-primed BMDMs (5×10^5^ cells) in 60-mm dish were transfected with 4μg of biotinylated-LPS (LPS EB-Biotin, Invivogen). After indicated times, cells were washed with cold PBS and lysed on ice with cell lysis buffer (1% Triton X-100, 50mM Tris/HCl pH 7.4, 150mM NaCl, 1mM EDTA and Complete Protease Inhibitor Cocktail (Roche)). Whole cell extracts (WCE) were collected after 14,500rpm centrifugation at 4°C for 15min, and aliquoted for input. Remaining WCE were incubated with Avidin beads (50:50 slurry, Pierce™ NeutrAvidin™ UltraLink™ Resin, Thermo Fisher Scientific) on a rocker at 4°C overnight. WCE-bead mixtures (Biotin pull-down) were washed five times with cell lysis buffer by centrifugation, and boiled for further SDS-PAGE and immunoblotting analysis.

#### Biotin labelling of cell surface proteins

BMDMs were seeded at 2.5×10^6^ cell/dish into 60-mm dish, primed with LPS, and transfected with LPS. After indicated times, surface proteins of cell were labelled with biotin (*34*). In brief, cells were washed twice with DBPS^+^ solution (Dulbecco’s Phosphate-Buffered Saline (Gibco) supplemented with 0.9mM CaCl_2_ and 0.49mM MgCl_2_, pH 7.4), incubated with 2.5mg/ml of biotin (EZ-Link™ Sulfo-NHS-LC-LC-Biotin, Thermo Fisher Scientific) in DPBS^+^ solution at 4°C for 30min, washed three times in 100mM glycine for 5min and further washed two times in 20mM glycine for 5min. Cells were then lysed with lysis buffer (1% Triton X-100, 50mM Tris/HCl pH 7.4, 150mM NaCl, 1mM EDTA and Complete Protease Inhibitor Cocktail (Roche)), and a portion of the whole cell extracts (WCE) aliquoted for input. The remaining WCE was incubated with Avidin beads (Pierce™ NeutrAvidin™ UltraLink™ Resin, Thermo Fisher Scientific) on a rocker at 4°C overnight. WCE-bead mixtures (Biotin pull-down) were washed six times with lysis buffer, and boiled for immunoblotting.

#### Cell fractionation

LPS-primed BMDMs (5×10^6^ cells/dish in 10-cm dish) were activated by LPS transfection as described above. At the indicated times, cells were separated into soluble and membrane fraction by hypotonic cell separation method. In brief, cells were detached by 0.5% EDTA solution and collected by 350x *g* centrifugation. After being washed twice with cold PBS, cells were incubated with hypotonic buffer (10mM Tris-HCl, pH 7.5) on ice for 10min, and homogenized by 25strokes of Dounce homogenizer (Sigma-Aldrich), and centrifuged 14,500rpm at 4°C for 15min. Supernatants were collected as the soluble fraction and the remaining insoluble pellets were washed with hypotonic buffer by 14,500 rpm centrifugation at 4°C for 15min. Supernatants were completely removed and the remaining insoluble pellets (insoluble membrane fraction) were incubated with sample lysis buffer (50mM Tris-HCl, pH 7.5, 150mM NaCl, and 1% Triton X-100) on ice for 30min. Soluble- and insoluble membrane-fractions were boiled for further immunoblotting analysis. For analysis of Native-PAGE, LPS-activated BMDMs were separated by hypotonic cell separation method as mentioned above with membrane protein extraction solution #6 (Profoldin, Hudson, MA) for membrane fraction. Each fraction was loaded into native gel (4-12% gradient). For efficient transfer to PVDF, the native gel was incubated with 0.3% SDS on rocker at RT for 1h (sodium dodecyl sulfate [SDS] soaking step) and immunoblotted with anti-Gsdmd antibody.

#### Gsdmd-N oligomerization and Co-IP

293T cells were seeded at 5×10^6^ cells into 10-cm dish, and transfected with 10μg of doxycycline-inducible plasmids containing Gsdmd-N or Gsdmd-N/RFP-Gbp3 or Gsdmd-N-GFP/RFP-Gbp3^S46N^ by TransIT-LT1 transfection reagent (Mirus Bio) according to manufacturer’s instruction. Cells were incubated with 1μg of doxycycline for 24h, washed and harvested with cold PBS. Soluble and insoluble membrane fractions were separated by the hypotonic cell fractionation method as described above. Each fraction was divided for either non-reducing (without 2-mercaptoethanol) and reducing conditions (with 2-mercaptoethanol and boiling). Samples were further analyzed by immunoblotting. To verify protein-protein interactions, soluble and insoluble membrane fractions were incubated with 1μg of anti-GFP antibody and protein A/G beads (pierce protein A/G agarose, Thermo Scientific) on a rocker at 4°C overnight. Sample-bead mixtures were washed six times with each fractionation buffer, and boiled for immunoblotting.

#### Co-immunoprecipitation

293T cell were seeded at 1×10^6^ cells/well into 6-well plates, and co-transfected with pMX-IP-HA-Gsdmd full-length, N-term Gsdmd^I105N^, N-term Gsdmd^L193D^, C-term and pCMV-Myc-Flag-Gbp3. After 24h, cells were lysed with SDS-containing cell lysis buffer (25mM Tris-HCl pH 7.4, 150mM NaCl, 5% Glycerol, 1% Triton X-100, and 0.1% SDS, protease inhibitor cocktail (Roche)) on ice. Whole cell lysates (WCL) were incubated with 1μg of anti-HA antibody (BioLegend) and protein A/G beads (pierce protein A/G agarose, Thermo Scientific) at 4°C overnight. Bead mixtures were washed 6times with cell lysis buffer, and boiled for immunoblotting.

#### Protein purification

Recombinant Flag/HA-tagged Gbp3 wild-type and its mutant proteins (K45A/S46A, S46N, D176N and R3A), Myc/Flag-tagged Gbp3, Myc-tagged Gbp2, Flag-tagged Caspase-11 were purified via a mammalian expression system. In brief, 293E cells were seeded at 1.6×10^7^ cells/dish into a 15-cm dish and transfected by 20μg of plasmid with LT-1 transfection reagent (Mirus Bio) according to manufacturer’s instructions, respectively. After 24h, cells were harvested by centrifugation and washed two times with cold PBS. Cell pellets were lysed with lysis buffer (50mM Tris-HCl, pH 7.4, 150mM NaCl, and 1% Triton X-100, protease inhibitor cocktail (Roche)) on ice, and cell lysates were loaded into equilibrated anti-Flag M2 affinity resin (Sigma-Aldrich) or anti-c-Myc Tag (9E10) Affinity Gel (BioLegend) according to column chromatography method of manufacturer’s instructions. Flag/HA-tagged proteins were eluted by competition with the 1x or 3x Flag peptides (Sigma-Aldrich), and Myc-tagged proteins were eluted by c-Myc peptide tag (BioLegend). Purified proteins were desalted into 50mM Tris-HCl pH 8.0, 100mM NaCl, 5mM MgCl_2_, 10% glycerol on an AKTA FPLC (GE Healthcare) and stored in aliquots at −80°C. Protein purity was confirmed by Coomassie blue staining (SimplyBlue™ SafeStain, Thermo Fisher), and protein concentration were determined by BCA assay (Pierce™ BCA Protein Assay Kit, Thermo Fisher). Recombinant 6x His/Flag-tagged Gsdmd full-length, N-term and C-term proteins were purified from bacterial expression system by *E. coli* (BL21DE3-RIPL).

#### *In vitro* protein interaction

Immunoprecipitation assay and Ni-NTA pull down assay were used for *in vitro* protein interaction studies (Interaction between Flag-Caspase-11 and Myc-Gbp2; Interaction between Myc-Gbp3 and 6xHis-Gsdmd full-length, N-term and C-term). For *In vitro* immunoprecipitation assay, recombinant proteins were incubated (1:1 ratio, each 200-300 ng of protein) in binding buffer (20mM HEPES and 0.5% Triton X-100) for 30min RT. Mixtures were incubated with 0.5μg of anti-Flag antibody or 0.5μg of anti-Myc antibody at 4°C for 2h on rocker, and further incubated with Protein A/G beads at 4°C for 2h. Protein-beads mixtures were washed 6times with binding buffer and boiled for immunoblotting. For Ni-NTA pull down assay, Purified 6x His/Flag-tagged Gsdmd full-length, N-term, C-term and Flag/HA-tagged Gbp3 were mixed (1:1 ratio, each 100-200ng of protein) in binding buffer (50mM NaH_2_PO_4_ pH8, 0.5M NaCl), incubated Ni-NTA agarose (Qiagen) at 4°C overnight. Protein-bead mixtures were washed 6times with binding buffer and boiled for immunoblotting.

#### Thin layer chromatography and size exclusion chromatography

GTPase activity of wild-type Gbp3 and its mutant proteins was analyzed via thin layer chromatography as described (Kim et al., 2011). In brief, the hydrolysis of α-[^32^P]GTP was carried out by using 5μM purified proteins (20mM HEPES pH7.5, 135mM NaCl, 5mM KCl, 1mM MgCl_2_) and assayed at 25°C. The reaction was started by addition of 100μM GTP, 10μCi α-[^32^P]GTP/mL and was quenched with 142mM EDTA after 30min. GTPase hydrolysis products GTP, GDP and GMP were separated by CEL 300 PEI plates (Sigma-Aldrich) with fluorescent indicator (UV254) using 750mM KH_2_PO_4_, pH3.5 as solvent and visualized by autoradiography. Oligomerization status of proteins was assayed by size exclusion chromatography (SEC) as described. In brief, self-assembly of Gbp3 wild-type and its mutant proteins were carried out using an analytical Superdex 200 (GE Healthcare) SEC column calibrated with gel-filtration standards (Bio-Rad). Proteins analyzed for their oligomeric status both in nucleotide-free (50mM Tris-HCl pH 8.0, 2mM MgCl_2_) and well as their nucleotide-bound status (50mM Tris-HCl pH8.0, 2mM MgCl_2_, 300μM GDP, 300μM AlCl_3_, 10mM NaF). Chromatographic protein size markers were used for generating a standard curve (fig. S8D) and Stokes radius derived from SEC according to La Verde *et al.* (*35*).

#### *In vivo* mouse sepsis model

For caspase-11-driven sepsis, mice (6-12 weeks old) were intraperitoneally injected with 4mg/kg of Poly(I:C) LMW (Invivogen). After 6-7h, mice were intraperitoneally injected again with 1mg/kg of LPS (O111:B4, Sigma-Aldrich) or equal volume of PBS as a vehicle control. Surface body temperature were determined by Infrared-Thermometer (IR-B152, Braintree Scientific) every 3h point between 0 to 24h, and every 12h point between 24 to 72h.

Survival rate was monitored every 12h up to 120h. Sera were collected at 18h point to measure serum IL-1β. Tissues were isolated from mice injected with poly (I:C) LMW and LPS intraperitoneally as described above. After 18h, mice were sacrificed and tissues were isolated, washed with cold PBS and homogenized with lysis buffer (1% Triton X-100, 50mM Tris/HCl pH 7.4, 150mM NaCl, 1mM EDTA and Complete Protease Inhibitor Cocktail (Roche)). Tissue homogenates were centrifugated 5 times at 14,500rpm at 4°C for 20min to remove debris, and boiled for immunoblotting. To analyze membrane Gsdmd-N *in vivo*, splenocytes were isolated at 18h post-LPS, passed through a 70μm cell strainer (Thermo Fisher), and single-cell suspensions washed twice with cold PBS. Red blood cells were removed by incubation of RBC lysis buffer (150mM NH_4_Cl, 10mM KHCO_3_, 0.1mM EDTA, pH 7.4) on ice for 5min. Splenocytes were counted and cytosolic- and membrane-fractions were separated via fractionation kit (Cell signaling). For Tlr4-driven sepsis, mice (6-12 weeks old) were intraperitoneally injected with 25mg/kg LPS (O111:B4, Sigma-Aldrich). Survival rate was monitored every 12h up to 240h.

#### Immunoblotting

Cell extracts, cell supernatants, tissue extracts and purified proteins were separated by SDS-PAGE gel electrophoresis (Bio-Rad), transferred to PVDF membranes (Bio-Rad), immunoblotted with the indicated antibodies (Table S5) and detected with Clarity ECL substrate (Bio-Rad) by a ChemiDoc™ MP Imaging System (Bio-Rad).

#### Statistics

Data were analyzed by GraphPad Prism 8.0 software. Statistical significance was determined by t-test (two-tailed) or one-way ANOVA (Dunnett’s multiple comparison tests) or two-way ANOVA (Multiple comparisons). All metrics are given as means ± standard deviation. p < 0.05 was considered statistically significant. Kaplan-Meier survival curve with Gehan-Breslow-Wilcoxon test were used for analyzing survival rate.

---

**Fig. S1.**
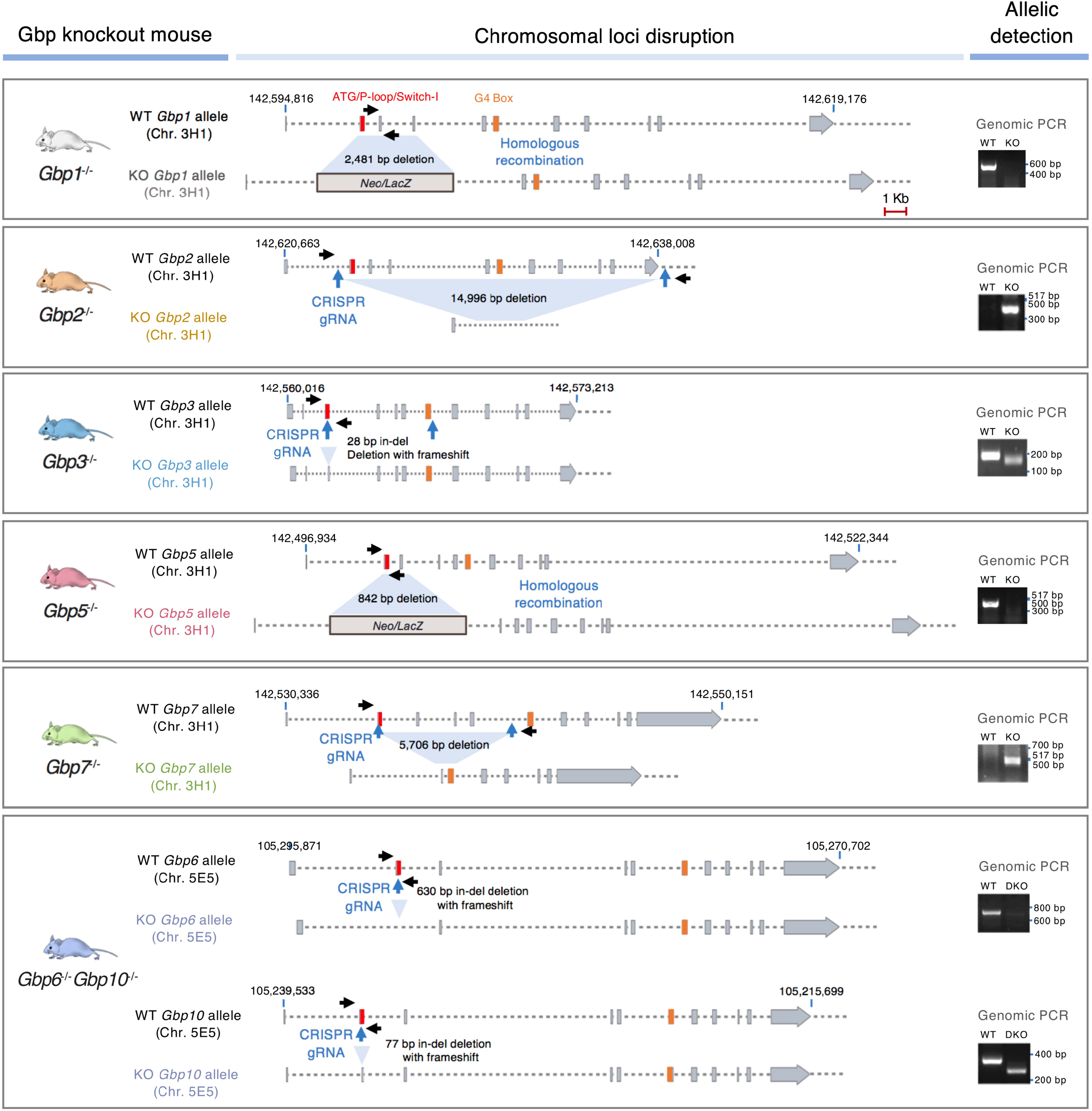
Gbp-deficient mice generated by CRISPR-Cas9 or homologous recombineering. Position of sgRNA insertions, deletions and frameshifts in murine *Gbp* loci to generate a panel of new knockout mice including *Gbp2^-^/^-^*, *Gbp3^-^/^-^*, *Gbp7^-^/^-^* and *Gbp6^-^/^-^Gbp10^-^/^-^* on chromosome 3H1 or 5E5. For *Gbp1^-^/^-^* and *Gbp5^-^/^-^* mice generated via homologous recombination, position of the *Neo^R^*/*LacZ* cassette is shown. Genomic PCR for allelic detection shown at right. All chromosomal alterations confirmed by genome sequencing.

**Fig. S2.**
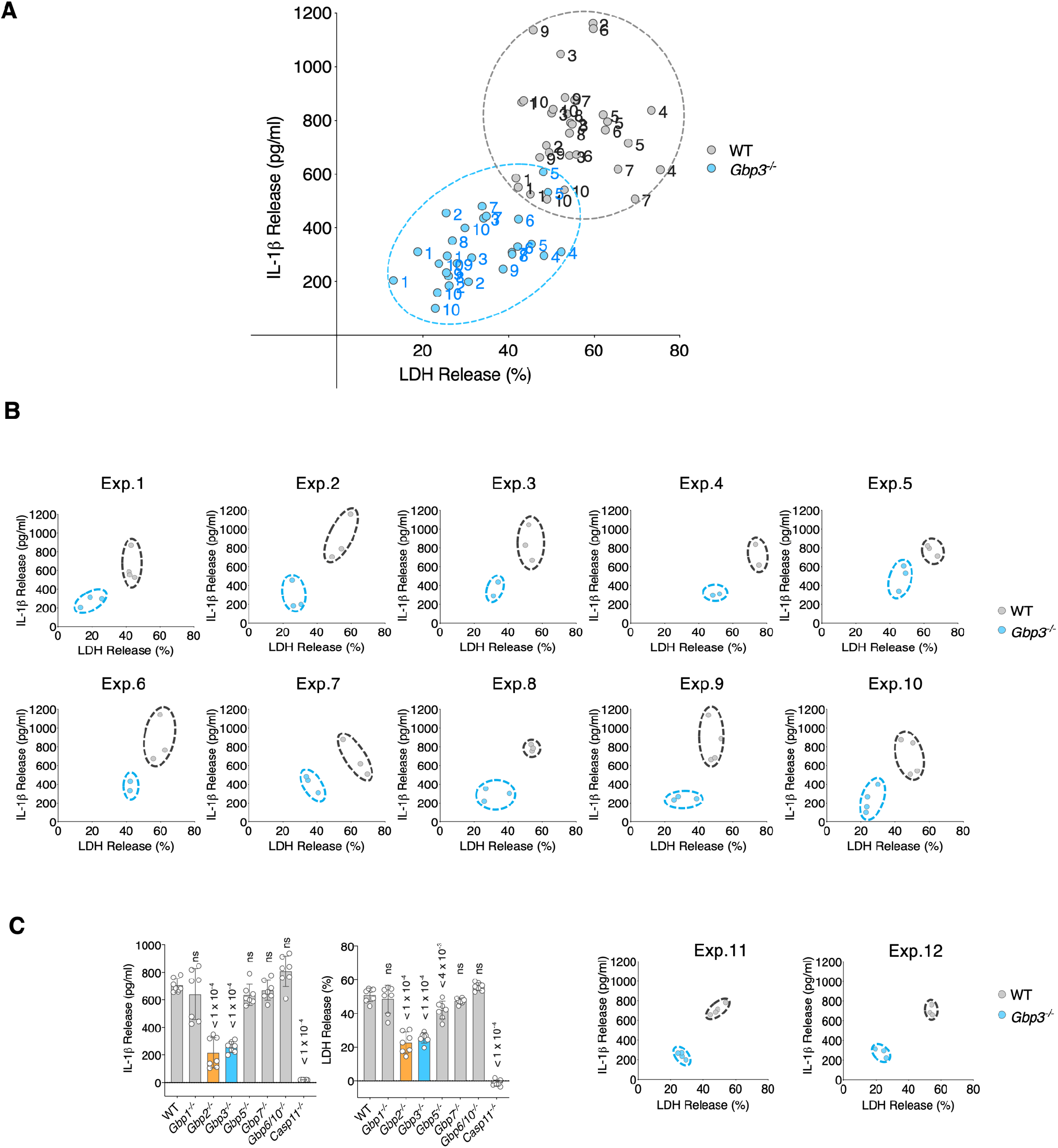
Defective iLPS-induced responses across multiple experiments using BMDMs from different *Gbp3^-^/^-^* mice. **(A).** An example of individual IL-1β and LDH measurements from 10 independent biological experiments (20 different mice) for *Gbp3^-^/^-^* versus WT BMDMs shown as a compiled scatter plot. Each individual experiment is numbered **(B).** The ten different biological experiments from the scatter plot showing duplicate, triplicate or quadruplicate samples measured for both genotypes. Defects in *Gbp3^-^/^-^* versus WT BMDMs are evident in each experiment **(C).** Experiments #11 and #12 conducted simultaneously alongside six other genotypes as shown Fig. 1C further underscore the importance of new *Gbp3^-^/^-^* mice revealing Gbp3 is important for iLPS responses.

**Fig. S3.**
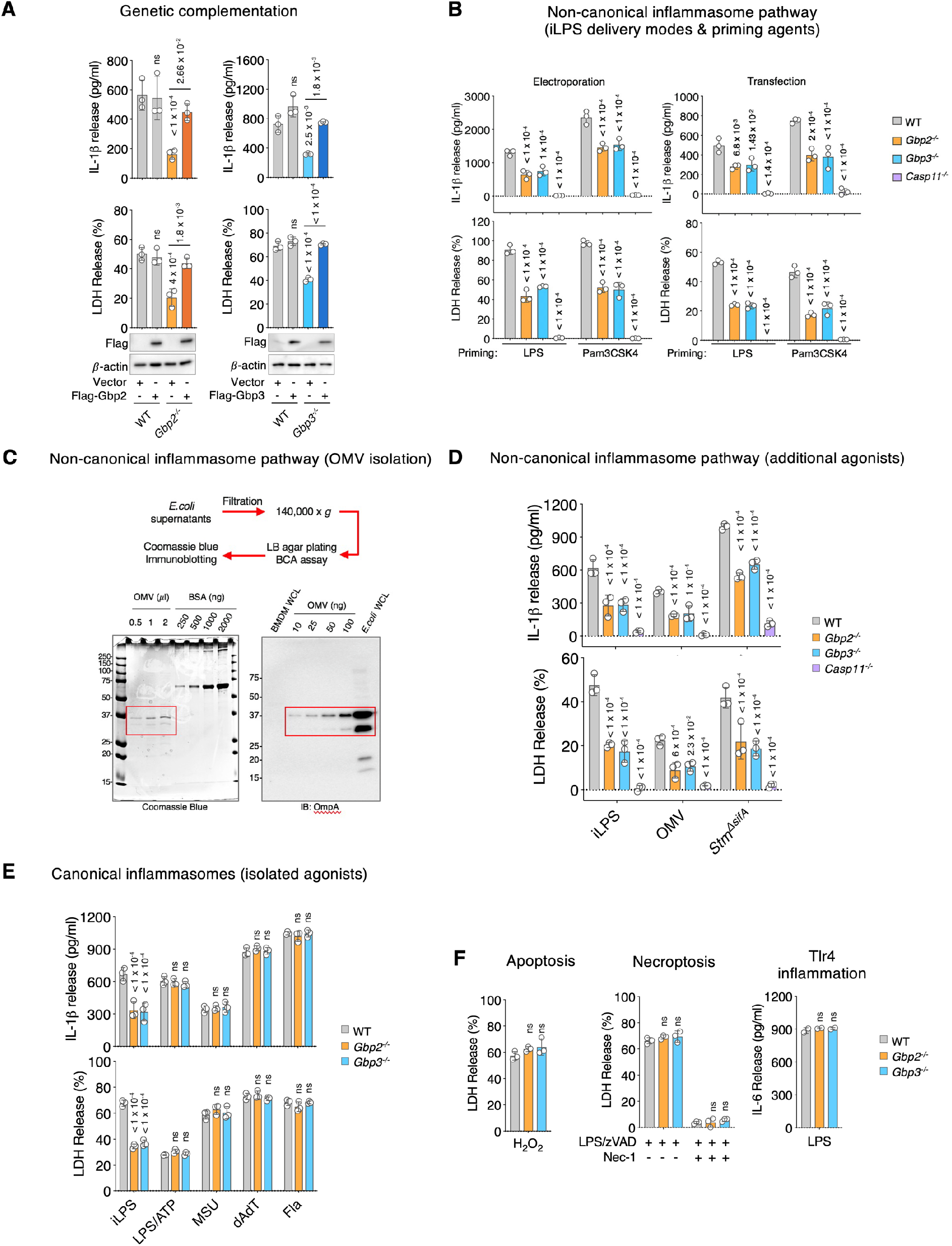
*Gbp2^-^/^-^* and *Gbp3^-^/^-^* BMDM responses to additional inflammasome, cytotoxic and immune stimuli. (**A**). Complementation of *Gbp2^-^/^-^* and *Gbp3^-^/^-^* BMDMs with Flag-Gbp2 or Flag/HA-tagged Gbp3 rescued defects in IL-1β and LDH release in response to iLPS*. P* values, one-way ANOVA with Turkey’s multiple comparison test versus WT control expressing empty vector. ns, not significant. Representative of 2 independent repeats. (B). Comparison of transfection and electroporation methods for introducing iLPS. Representative of 3 independent experiments. (**C**). Coomassie stain and immunoblot of outer membrane vesicle (OMV) fractions prepared from *E. coli* as shown in the schematic above. (**D**). Cytokine and pyroptotic defects in *Gbp2^-^/^-^* and *Gbp3^-^/^-^* BMDMs exposed to additional caspase-11 activators including *E.coli*-derived OMVs and SifA-deficient *S. typhimurium*. *P* values, one-way ANOVA with Dunnett’s multiple comparison test versus WT controls. Representative of 3 independent repeats. (**E**). Intact cytokine and pyroptotic responses in *Gbp2^-^/^-^* and *Gbp3^-^/^-^* BMDMs exposed to isolated activators of canonical inflammasomes compared with iLPS. Agonists used for Nlrp3 (LPS/ATP; MSU), Aim2 (dA:dT) and Nlrc4 inflammasomes (monomeric flagellin). *P* values, one-way ANOVA with Dunnett’s multiple comparison test versus WT controls. ns, not significant. Representative of 3 independent repeats. (**F**). Intact apoptotic, necroptotic and LPS-induced Tlr4 cytokine responses in *Gbp2^-^/^-^* and *Gbp3^-^/^-^* BMDMs. Apoptotic cell death in response to H_2_O_2_, necroptotic cell death in response to LPS/zVAD with the inhibitor necrostatin used as a control, and IL-6 release in response to LPS. ns, not significant. Representative of 2 independent repeats.

**Fig. S4.**
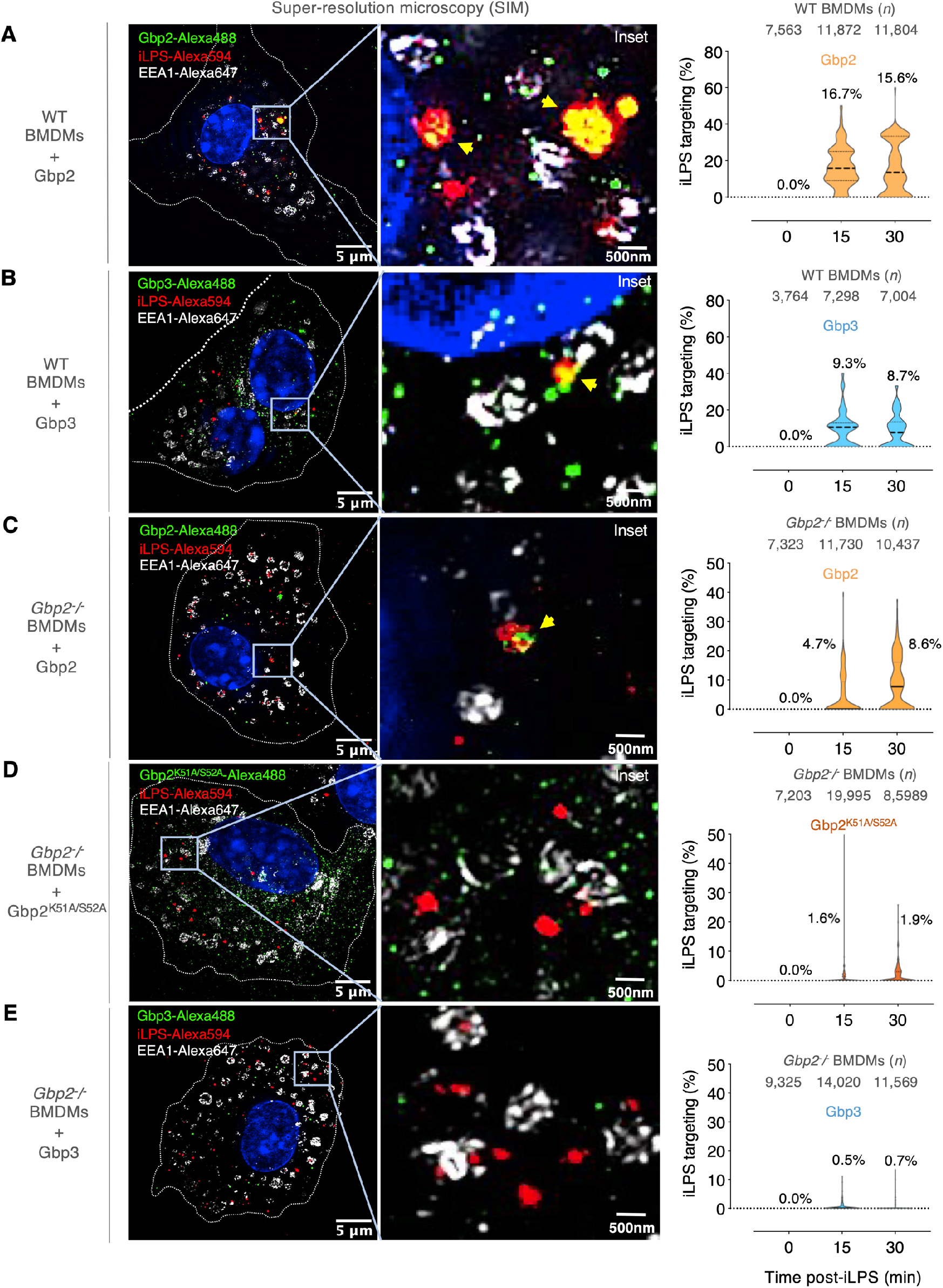
GTPase-active Gbp2 targets iLPS upstream of caspase-11 detection in primary BMDMs. Super-resolution microscopy (SIM) of BMDMs from WT mice expressing **(A)** Flag-Gbp2 or **(B)** Flag-HA-Gbp3. *Gbp2^-^/^-^* BMDMs expressed either **(C)** YFP-Gbp2; **(D)** YFP-Gbp2^K51A/S52A^ (GTPase mutant) or **(E)** YFP-Gbp3. Images captured 30 min after LPS uptake at peak Gbp colocalization (yellow arrows). iLPS detected via Alexa-594 and the early endosomal Rab5a effector, EEA1, via Alexa647. Large-scale unbiased computational enumeration (InCell) of Gbp and iLPS co-localization in violin plots. Number (*n*) of BMDMs analyzed at each time point.

**Fig. S5.**
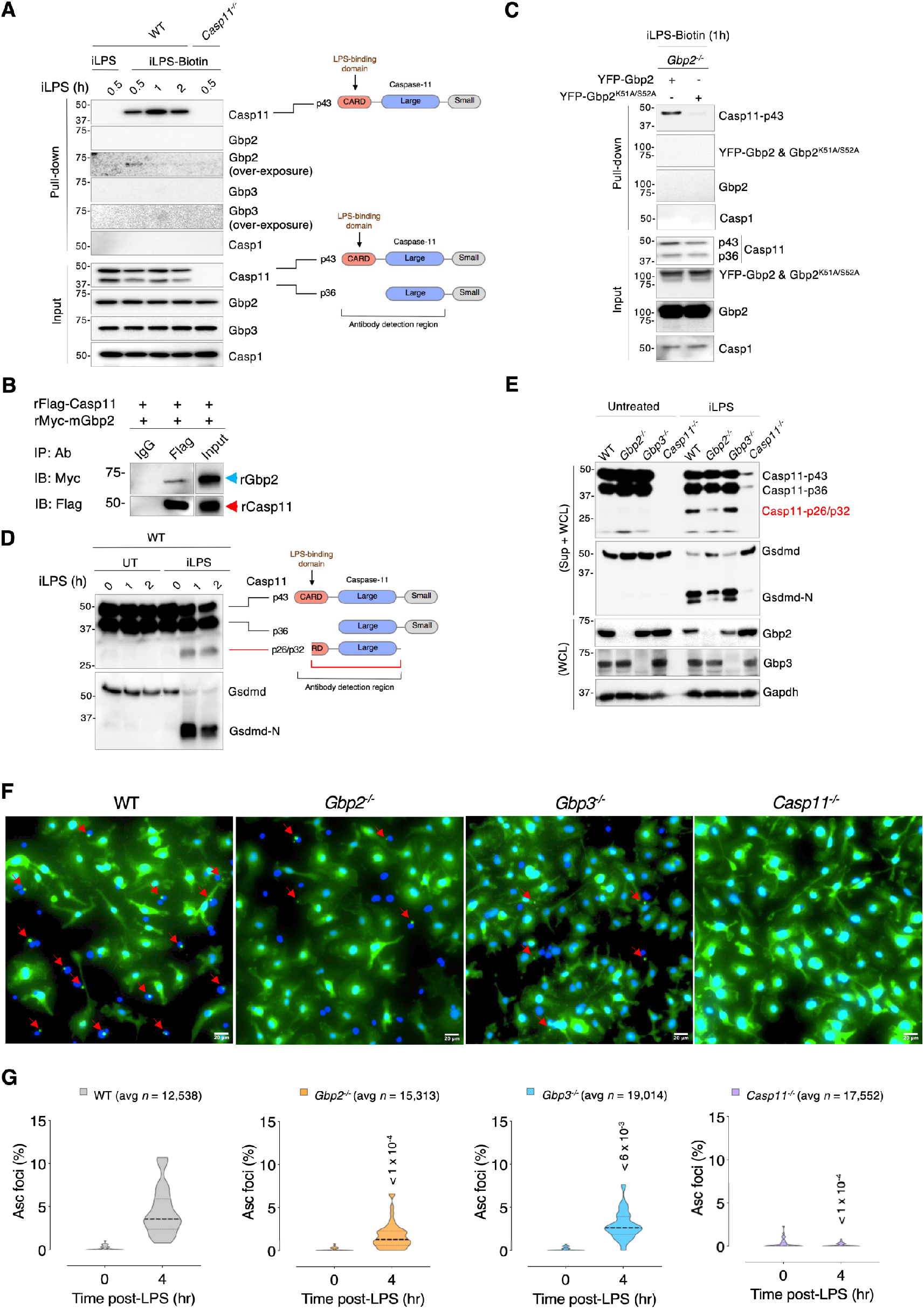
Gbp2 is required iLPS detection by caspase-11 to trigger downstream events. **(A).** Time course of iLPS-biotin binding to the endogenous CARD-containing caspase-11 p43 fragment in WT BMDMs. *Casp11^-^/^-^* BMDMs served as a negative control. Over-exposure of Gbp2 and Gbp3 immunoblotting showed weak binding by Gbp2 but not Gbp3 to iLPS-biotin as well. Caspase-11 antibody detection region shown at right. **(B).** Direct interaction of recombinant Myc-Gbp2 by recombinant Flag-caspase-11 in cell-free pulldown using anti-Flag antibody with IgG isotype control. **(C).** Ligand co-capture at peak 60 min post-uptake in *Gbp2^-^/^-^* BMDMs complemented with YFP-Gbp2 or YFP-Gbp2^K51A/S52A^. One of two replicates shown. **(D).** Time course of iLPS-induced caspase-11 and Gsdmd cleavage as depicted by p26 caspas11 fragment and ∼30kDa Gsdmd-N fragment in immunoblot. **(E).** Pronounced defects in caspase-11 autoactivation and Gsdmd cleavage in *Gbp2^-^/^-^* BMDMs shown in extended blots. **(F).** Formation of endogenous Asc foci (red arrows) stained with anti-Asc antibody in BMDMs from WT, *Gbp2^-^/^-^*, *Gbp3^-^/^-^* and *Casp11^-^/^-^* mice. Representative of three replicates. **(G).** Large-scale, unbiased enumeration of endogenous Asc foci in BMDMs from WT, *Gbp2^-^/^-^*, *Gbp3^-^/^-^* and *Casp11^-^/^-^* mice.

**Fig. S6.**
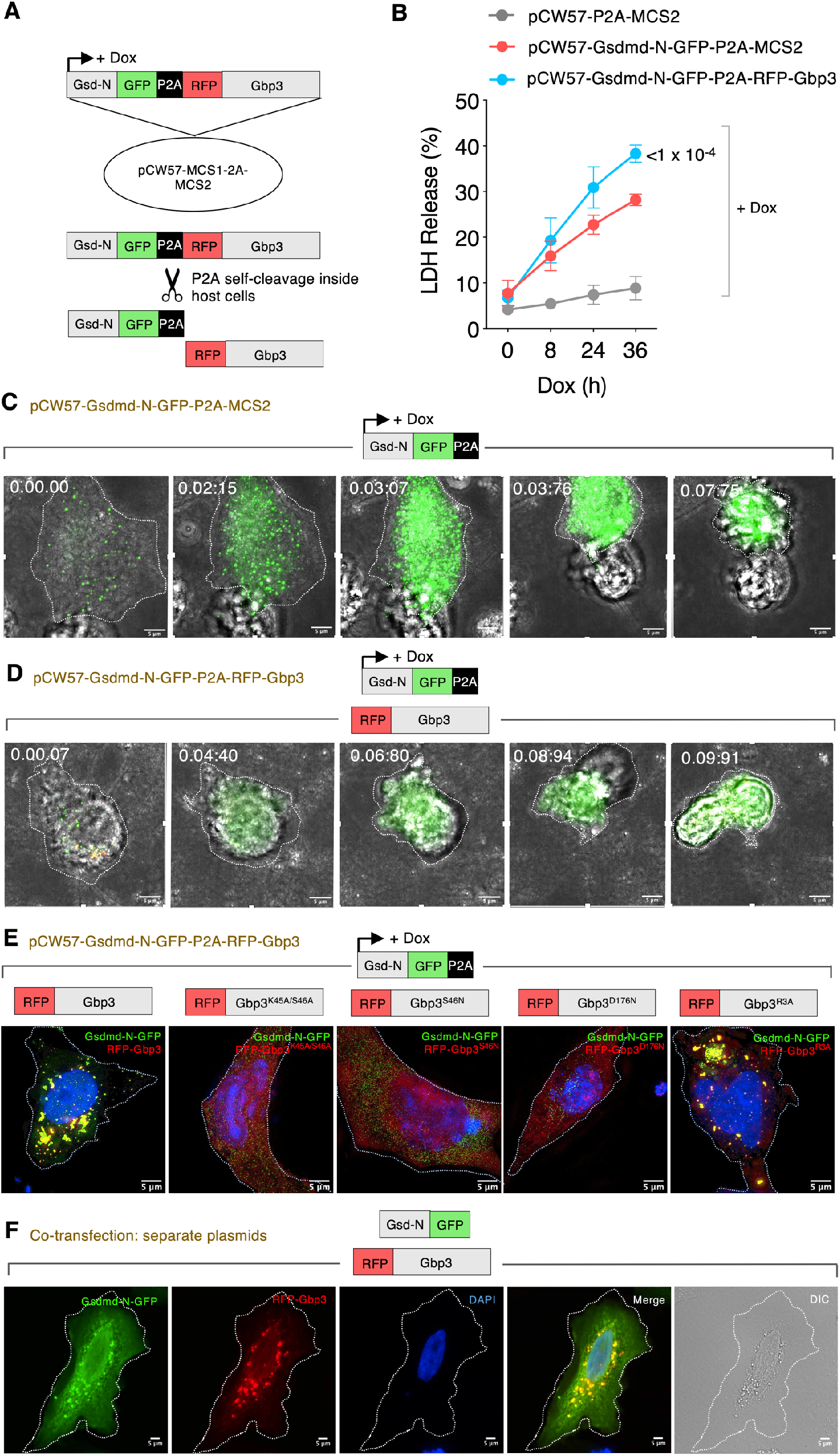
Development of a bipartite system for functional Gbp3 and Gsdmd-N expression. **(A).** A doxycycline (Dox)-inducible lentiviral bipartite P2A (self-cleaving peptide) expression system (pCW57-MCS1-2A-MCS2) to co-regulate Gsdmd-N and Gbp3-driven cytotoxicity. Gsdmd-N-GFP and RFP-Gbp3 produced within the same cell are separately released after P2A peptide self-cleavage. No uncleaved Gsdmdm-N-P2A-Gbp3 was detected in immunoblot. **(B).** Functional test of the bipartite system in HeLa cells. Addition of doxycycline (+ Dox) induced pyroptosis measured by LDH release in lentiviral transduced cells expressing pCW57-Gsdmd-N-GFP-2A-MSC2. Cytotoxicity was significantly increased when RFP-Gbp3 was co-expressed together with Gsdmd-N using pCW57-Gsdmd-N-GFP-2A-RFP-Gbp3. No increase in LDH release was observed without doxycycline (-Dox). **(C).** Live cell imaging of pyroptosis due to doxycycline-driven pCW57-Gsdmd-N-GFP-2A-MSC2 expression. **(D).** Live cell imaging of pyroptosis due to doxycycline-driven pCW57-Gsdmd-N-GFP-2A-RFP-Gbp3 expression. **(E).** Super-resolution (SIM) imaging of pCW57-Gsdmd-N-GFP-P2A-RFP-Gbp3 plasmids harboring different Gbp3 mutants in HeLa cells for comparison with S6F below. Loss of colocalization for RFP-Gbp3^K45A/S46A^, RFP-Gbp3^S46N^ and RFP-Gbp3^D176N^ with Gsdmd-N-GFP-P2A demonstrate P2A is fully active. Thus, convergence of RFP-Gbp3 or RFP-Gbp3^R3A^ with Gsdmd-N-GFP-P2A is not due to uncleaved bipartite expression. **(F).** 3-color imaging plus DIC reveal separately transfected plasmids encoding Gsdmd-N-GFP and RFP-Gbp3 strongly colocalize like pCW57-Gsdmd-N-GFP-P2A-RFP-Gbp3, further reinforcing this relationship inside mammalian cells.

**Fig. S7.**
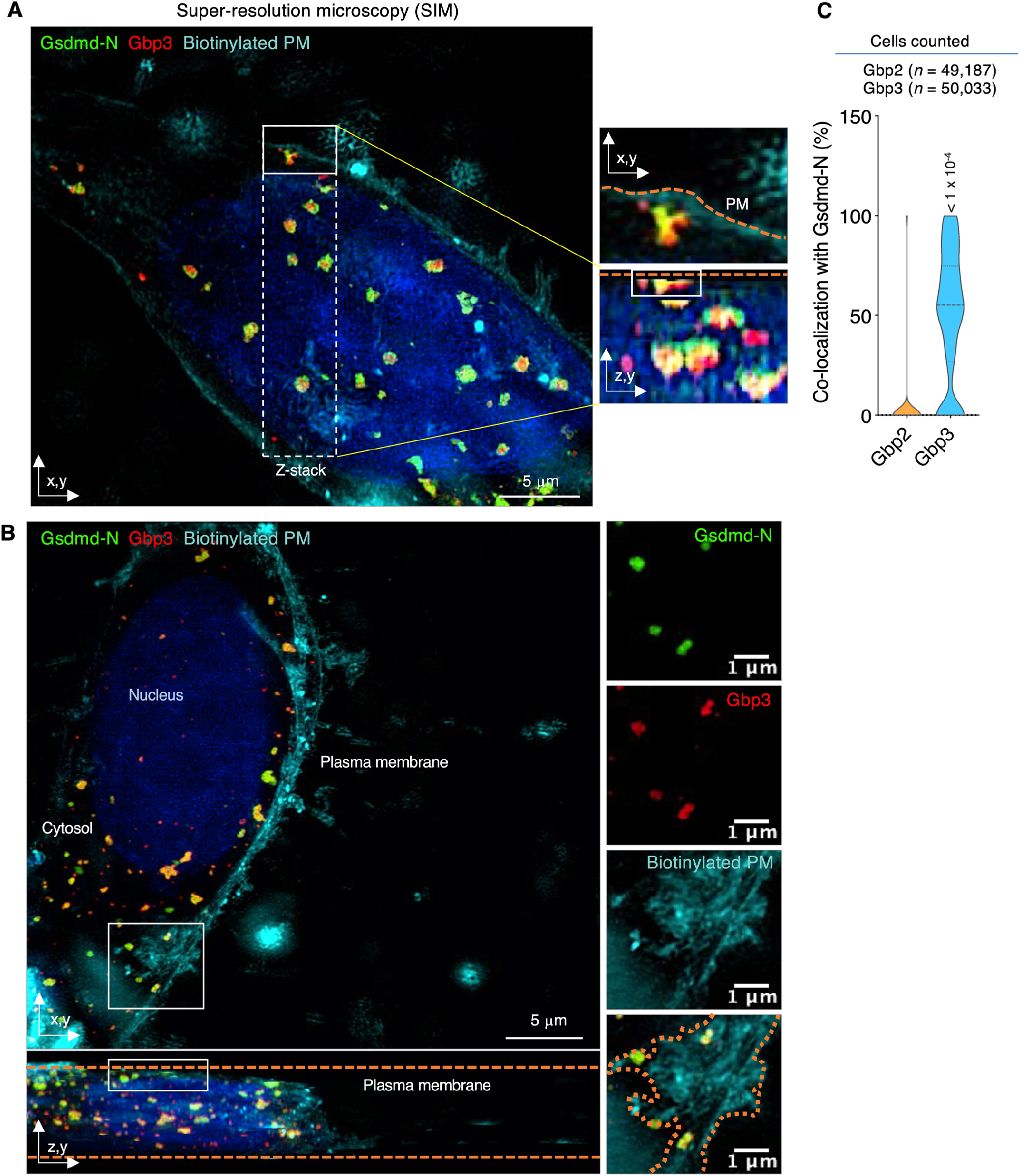
Gbp3 and Gsdmd-N generate large colocalized structures inside mammalian cells. **(A).** Four-color SIM imaging of RFP-Gbp3 and Gsdmd-N-GFP in transfected HeLa cells. Plasma membrane labeling used biotin-Cy5 and shown in z,y planar view (below). DAPI was used to delineate the nucleus. **(B).** Individual SIM channels for Gsdmd-N, Gbp3 and biotinylated plasma membrane along with overlay containing the DAPI-labelled nucleus from the insets in a,b. Dotted lines depict z-plane boundaries for the cell along with plasma membrane edge. **(C).** Violin plots of Gbp2 or Gbp3 colocalization with Gsdmd-N identifed via unbaised computational sampling. One-way ANOVA with Dunnett’s multiple comparison test between the two Gbp-labelled groups.

**Fig. S8.**
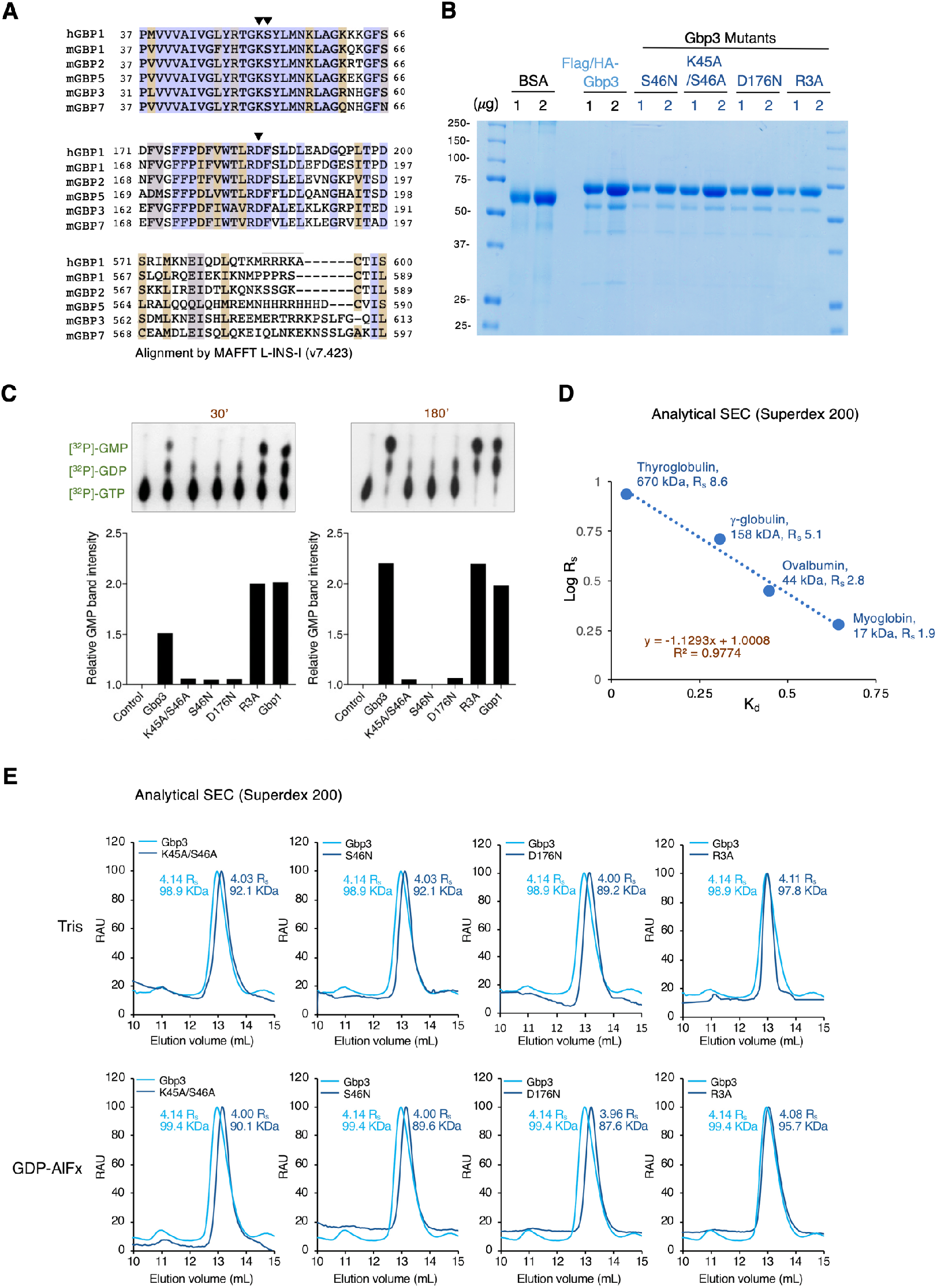
Gbp3 produces GMP and forms elongated conformers in size exclusion chromatography. **(A).** Sequence alignment of Gbp3 and selected murine Gbps with human GBP1 by MAFFT analysis depict residues needed for G-domain catalytic activity and assembly-driven GMP production. **(B).** Coomassie stain of recombinant Flag-HA-Gbp3 and its mutants isolated from human 293E cells to ensure eukaryotic post-translational modification. Protein loading shown with a BSA control. **(C).** GTP hydrolysis assay of recombinant mouse Gbp3 and its mutants isolated from human 293E cells at 30 and 180 min after addition of [^32^P]-labelled substrate. Thin-layer chromatography shown above with relative GMP band intensity quantified below. One of three replicates shown. **(D).** Size exclusion chromatography (SEC) standard curve using proteins of known globular size with the base logarithm of Stokes radius (Log R_s_) plotted against their experimentally determined solute behavior (distribution coefficient, Kd) in Superdex 200 columns. **(E).** SEC of recombinant mouse Gbp3 and its mutants in the absence (Tris buffer) or presence of the transition-state analogue, GDP plus aluminum fluoride (AIFx). Catalytically active Flag-HA-Gbp3 and Flag-HA-Gbp3^R3A^ are extended conformers (∼10kDa larger) than inactive Flag-HA-Gbp3^K45A/S46A^, Flag-HA-Gbp3^S46N^ and Flag-HA-Gbp3^D176N^.

**Fig. S9.**
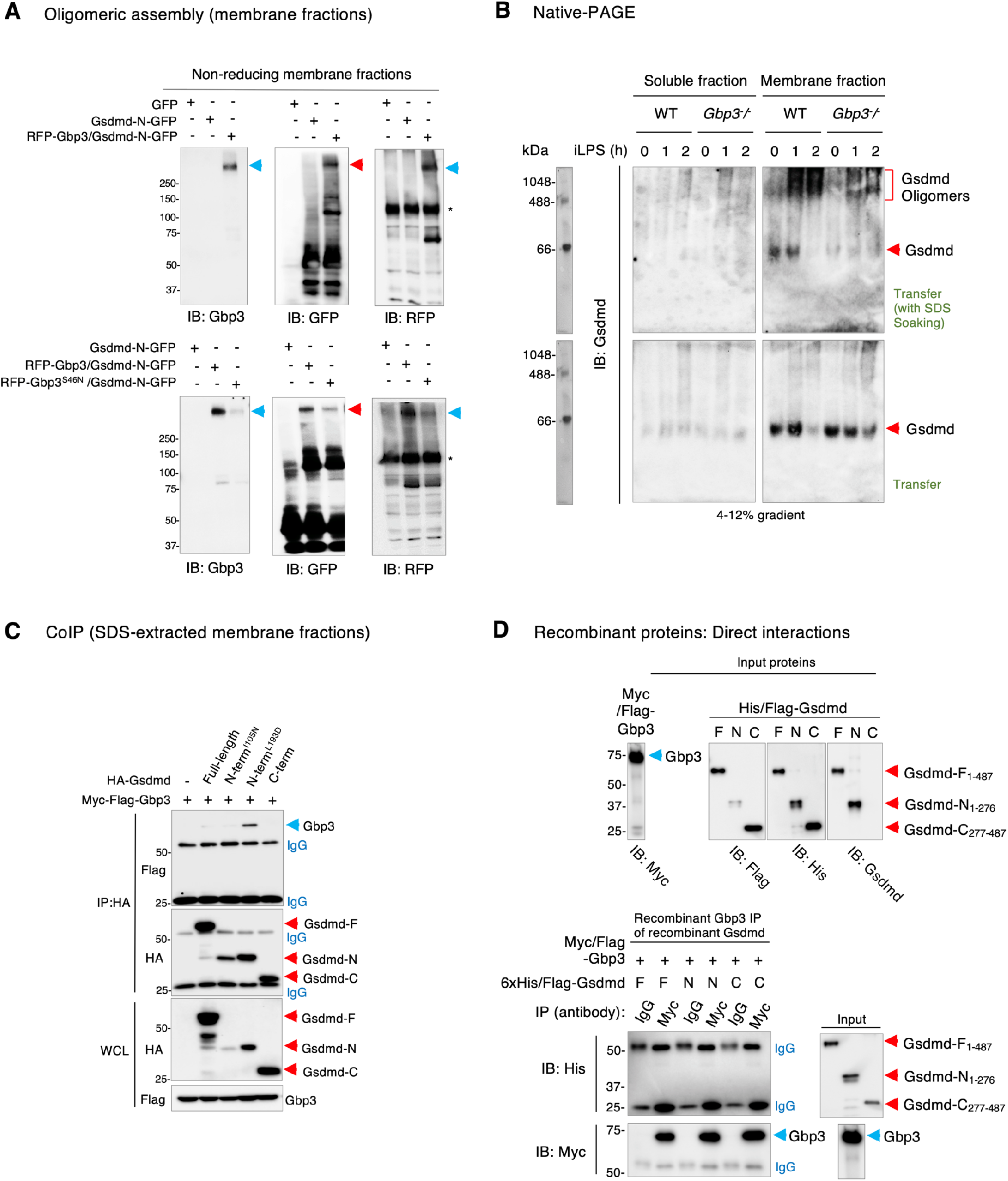
Catalytically- ctive Gbp3 interacts with Gsdmd-N to promote membrane complexes. **(A).** Assembly of large >250kDa complexes (arrows) of Gbp3 or Gsdmd-N in membrane fractions under non-reducing conditions (top). Substitution of RFP-Gbp3 with inactive RFP-Gbp3^S46N^ reduced these complexes (bottom). Antibodies used for immunoblot given below. Asterisk, non-specific band. One of two replicates shown. **(B).** Native PAGE of soluble and membrane fractions from WT and *Gbp3^-^/^-^* BMDMs treated with iLPS for 2 hours. Shift of native Gsdmd (arrows) into oligomeric structures after iLPS exposure which is largely lost in *Gbp3^-^/^-^* BMDMs (bracket). Soaking membranes in SDS during transfer enabled higher molecular weight species to be detected by anti-Gsdmd antibody. Apparent size marker shown (left). **(C).** Interaction of Myc-Flag-Gbp3 with hemagglutinin (HA)-Gsdmd variants (full-length, N- and C-terminal constructs) in co-immunoprecipitation assays of transfected 293T cells. IgG cross-reactivity for IP and IB demarcated. WCL, whole cell lysate. One of three replicates shown. **(D).** Pulldown assay for direct interactions between Gbp3 and Gsdmd. (Top) Recombinant input proteins for this assay immunoblotting with different antibodies to detect tags and intrinsic protein epitopes. Gsdmd fragment sizes shown on left. (Bottom) Recombinant coIP and Nickel-His_6_ column pulldown shows no increase of bidirectional retrieval versus IgG alone or non-specific background binding.

**Fig S10.**
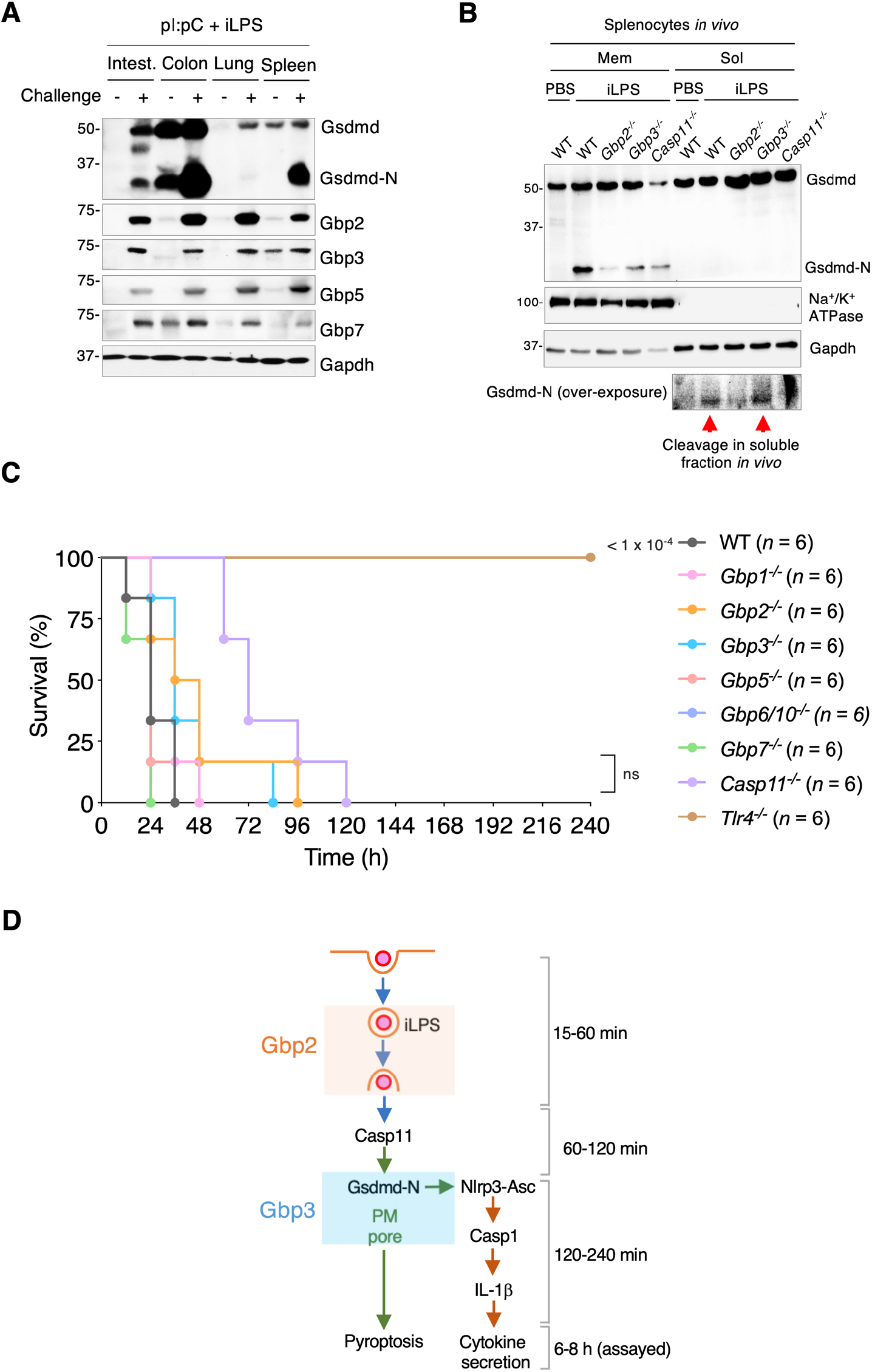
*Gbp2^-^/^-^* and *Gbp3^-^/^-^* mice succumb to Tlr4-dependent, caspase-11-independent sepsis. **(A).** Immunblot of native Gsdmd in homogenized tissue samples from pI:pC/iLPS-treated WT mice taken at 18 hr after challenge. Expression of Gbp2, Gbp3, Gbp5 and Gbp7 were also probed in these tissues. Asterisk, non-specific band. Based on the amount of cleaved Gsdmd-N produced specifically in response to iLPS, spleen (red box) was chosen as the source of material for membrane fractionation in Fig. 4d. **(B).** Gsdmd-N in membrane versus soluble fractions of splenocytes taken from WT and knoclout mice during caspase-11-dependent sepsis. Over-exposure of Gsdmd-N in soluble fractions reveal intact cleavage in splenocytes from WT and *Gbp3^-^/^-^* (arrows) but not *Gbp2^-^/^-^* and *Casp11^-^/^-^* mice as predicted. **(C).** LPS challenge across 9 genotypes with 25 mg/kg LPS (O111:B4). *Tlr4^-^/^-^* mice were included as a positive control and *Casp11^-^/^-^* mice as a negative control. *P* values for each knockout group shown versus wild-type C57BL6N mice using the Gehan-Breslow-Wilcoxon test. *n*, number of mice per group. **(D).** Co-operative model depicting the sequential steps at which Gbp2 and Gbp3 were found to impact the non-canonical inflammasome pathway. An approximate temporal scale is depicted for these events based on experiments described herein.

**Table S1.** Plasmids used in this study.

| Clone name | Source | Note |
| --- | --- | --- |
| pcDNA3.1-hygro-Flag/HA-mGbp3 | This study |  |
| pcDNA3.1-hygro-Myc/Flag-mGbp3 | This study |  |
| pCMV-VSV-G | Addgene | 8454 |
| pCMV-Flag-Caspase11 | Addgene | 21145 |
| pCMV-myc-mGbp1 | This study |  |
| pCMV-myc-mGbp2 | This study |  |
| pCMV-myc-mGbp3 | This study |  |
| pCW57-MCS1-2A-MCS2 | Addgene | 71782 |
| pCW57-MCS1-2A-MCS2-Gsdmd-N-term-GFP | This study |  |
| pCW57-MCS1-2A-MCS2-Gsdmd-N-term-GFP/RFP-mGbp2 | This study |  |
| pCW57-MCS1-2A-MCS2-Gsdmd-N-term-GFP/RFP-mGbp3 | This study |  |
| pCW57-MCS1-2A-MCS2-Gsdmd-N-term-GFP/RFP-mGbp3<br>K45A/S46A | This study |  |
| pCW57-MCS1-2A-MCS2-Gsdmd-N-term-GFP/RFP-mGbp3<br>S46N | This study |  |
| pCW57-MCS1-2A-MCS2-Gsdmd-N-term-GFP/RFP-mGbp3<br>D176N | This study |  |
| pCW57-MCS1-2A-MCS2-Gsdmd-N-term-GFP/RFP-mGbp3<sup>R3A</sup> | This study |  |
| pEGFP-c1 | Clontech | Empty vector |
| pEGFP-c1-mGbp3 | This study |  |
| pERPP-c1-mGbp2 | This study |  |
| pERFP-c1-mGbp3 | This study |  |
| pET28a-Flag-mGsdmd full-length | This study |  |
| pET28a-Flag-mGsdmd N-term | This study |  |
| pET28a-Flag-mGsdmd C-term | This study |  |
| pGag/Pol | Addgene | 14887 |
| Gsdmd (NM_026960) Mouse Tagged ORF Clone | Origene | MR207809 |
| pMXs-IRES-Puro | Clontech | Empty vector |
| pMXs-IRES-Puro-Flag-mGbp2 | This study |  |
| pMXs-IRES-Puro-Flag/HA-mGbp3 | This study |  |
| pMXs-IRES-Puro-Flag/HA-mGbp3<sup>K45A/S46A</sup> | This study |  |
| pMXs-IRES-Puro-Flag/HA-mGbp3<sup>S48N</sup> | This study |  |
| pMXs-IRES-Puro-Flag/HA-mGbp3<sup>D176N</sup> | This study |  |
| pMXs-IRES-Puro-Flag/HA-mGbp3<sup>R3A</sup> | This study |  |
| pMXs-IRES-Puro-GFP | This study |  |
| pMXs-IRES-Puro-YFP-mGbp2 | Shenoy *et al.*<br>(2012) |  |
| pMXs-IRES-Puro-YFP-mGbp2<sup>K51A/S52A</sup> | Shenoy *et al.*<br>(2012) |  |
| pMXs-IRES-Puro-HA-mGsdmd full-length | This study |  |
| pMXs-IRES-Puro-HA-mGsdmd N-term<sup>I105N</sup> | This study |  |
| pMXs-IRES-Puro-HA-mGsdmd N-term<sup>L193D</sup> | This study |  |
| pMXs-IRES-Puro-HA-mGsdmd C-term | This study |  |

**Table S2.**
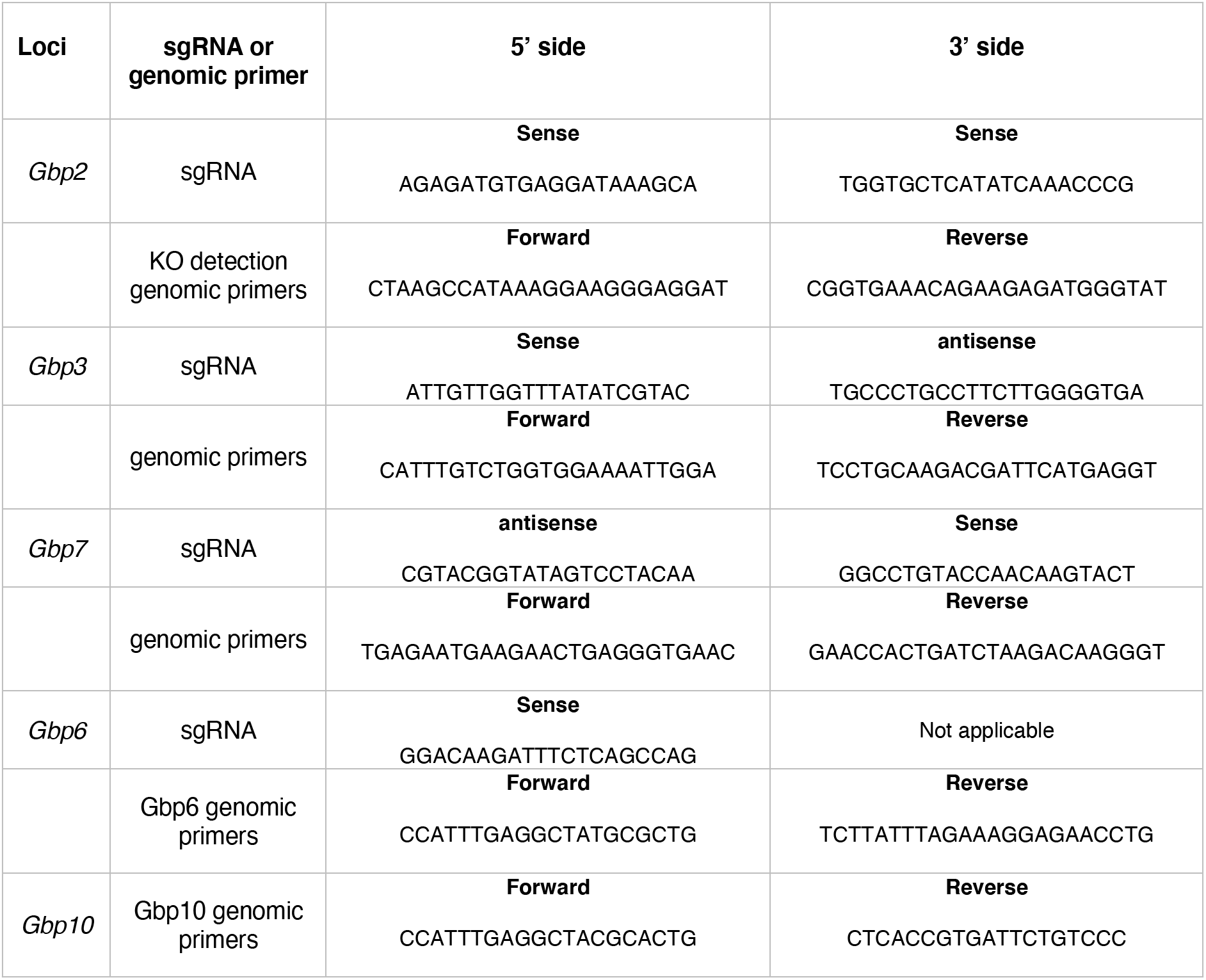
Guide sequences and genomic PCR primers for CRISPR-Cas gene targeting in mice.

**Table S3.**
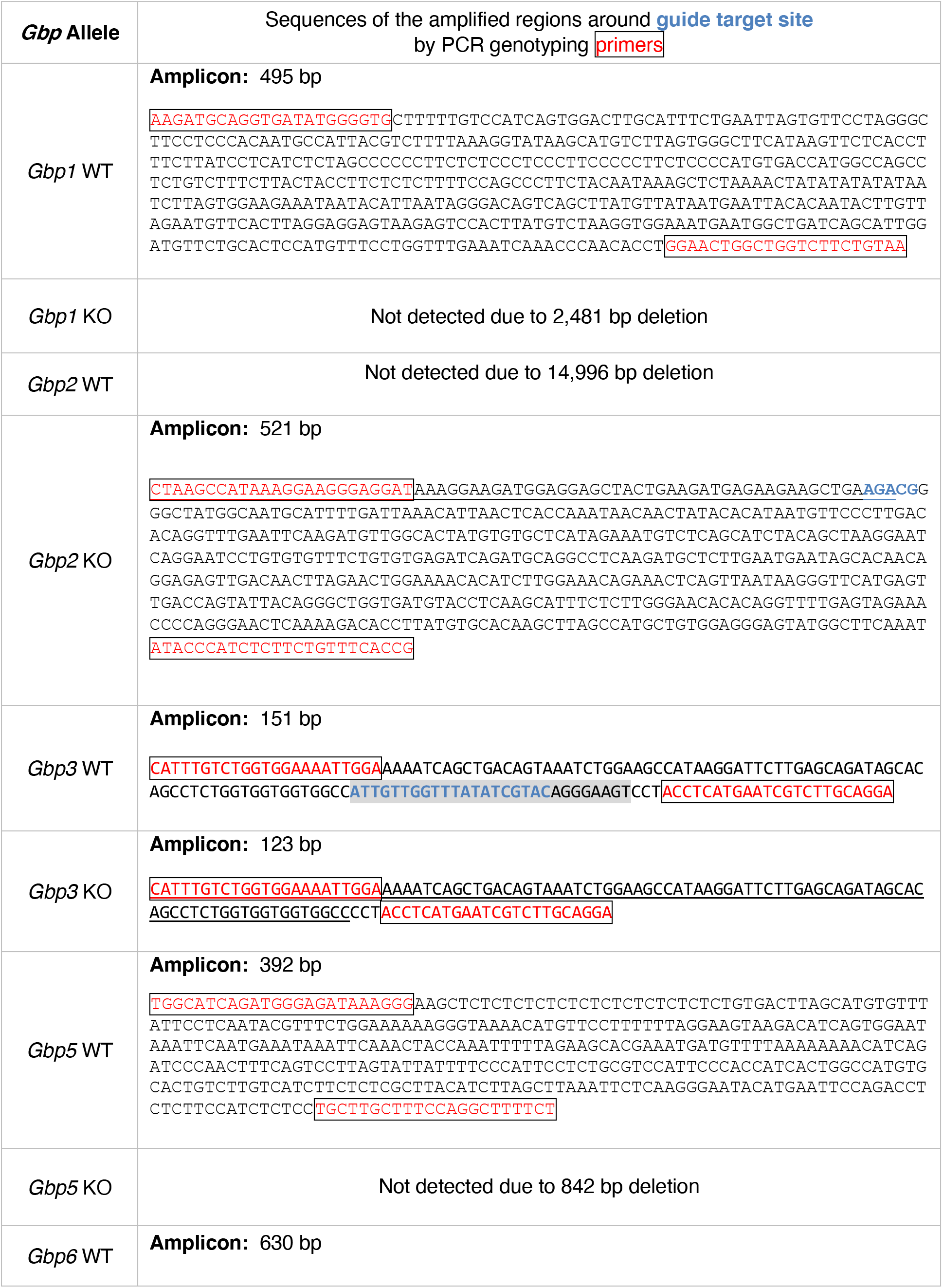
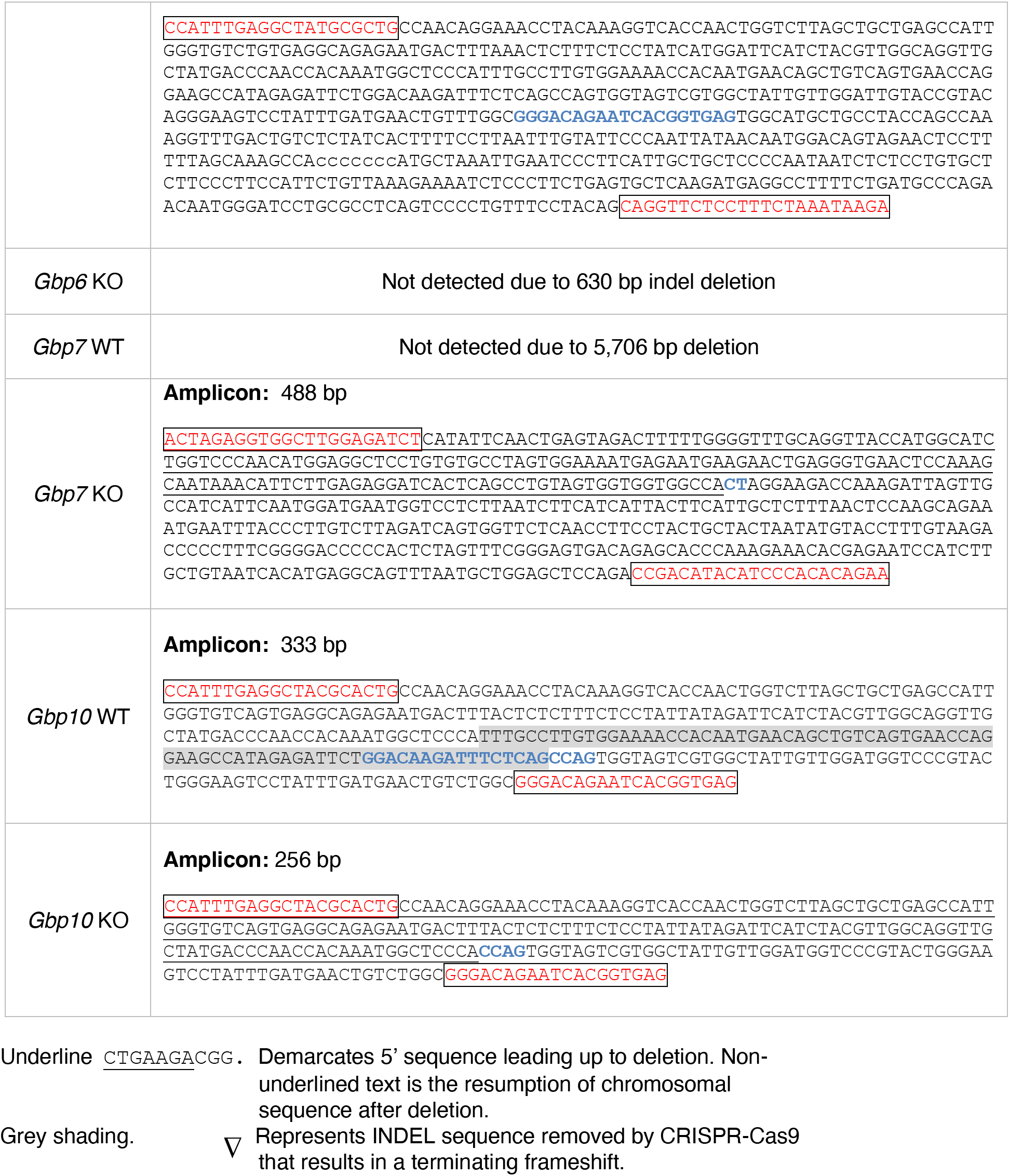
Amplified chromosomal regions from PCR genotyping of Gbp KO mice.

**Table S4.**
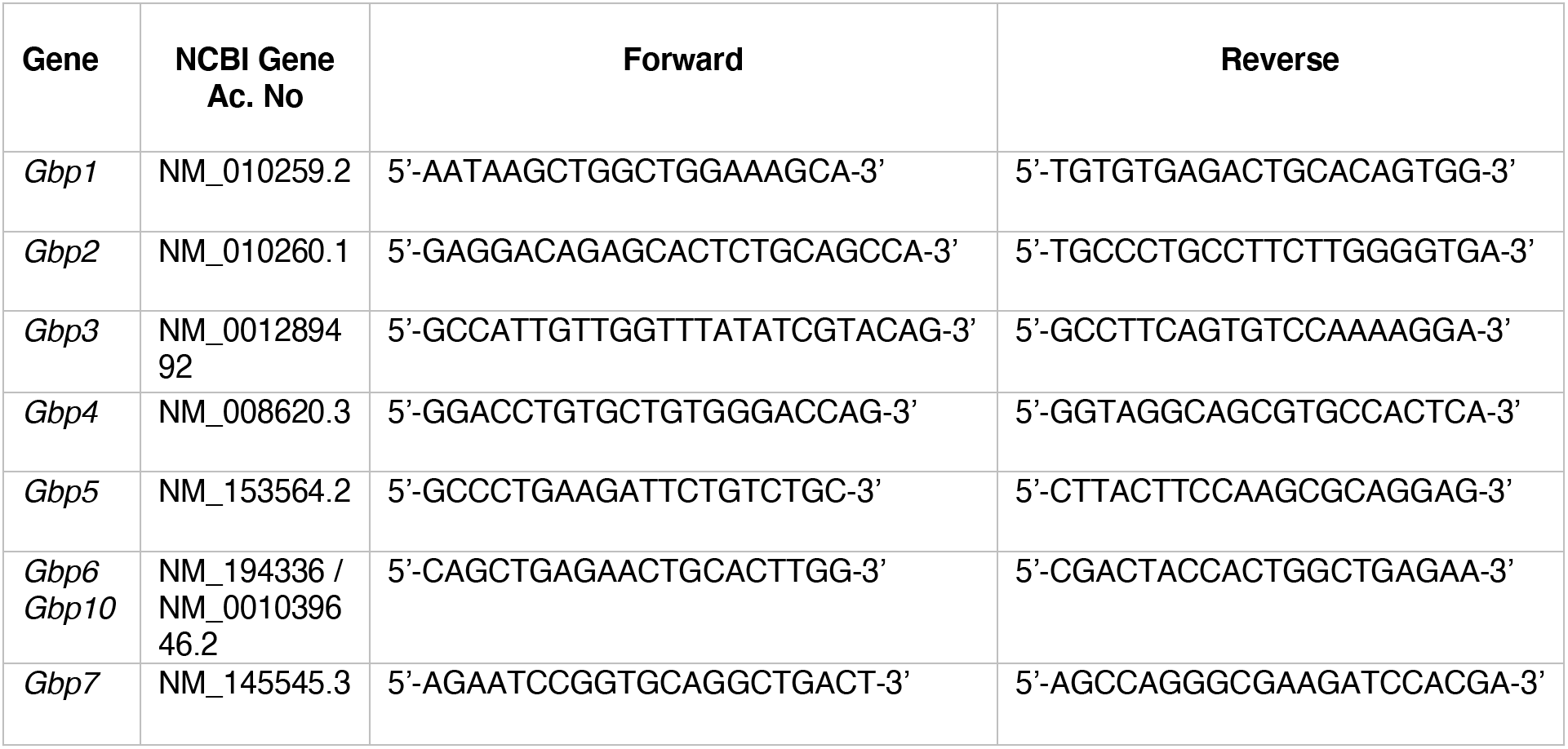
Primer sets for semi-quantitative RT-PCR.

**Table S5.** Antibodies used in this study.

| Antigen | Species<br>IgG/IgM | Application<br>(Dilution) | Product No. | Source | Note |
| --- | --- | --- | --- | --- | --- |
| ASC/TMS1 | Rabbit | IB (1:1000),<br>IF (1:500) | 67824 | Cell Signaling | D2W8U |
| Actin | Rabbit | IB (1:5000) | A5060 | Sigma-Aldrich | 20-33 |
| Caspase-1 | Mouse | IB (1:1000) | AG-20B-0042-C100 | Adipogen | p20, Casper-1 |
| Caspase-11 | Rat | IB (1:1000) | 14-9935-82<br>NB120-1-454 | eBioscience<br>Novus<br>Biologicals | 17D9 |
| Donkey anti-Goat IgG-HRP | Goat | IB (1:1000) | PA1-28664 | Thermo Fisher |  |
| DsRed | Mouse | IB (1:5000) | sc-390909 | Santa Cruz<br>Biotechnology | E-8 |
| EEA1 antibody | Rabbit | IF (1:1000) | 2411 | Cell Signaling |  |
| FLAG (M2) | Mouse | IB (1:10000),<br>IF (1:1000) | F1804 | Sigma-Aldrich |  |
| GAPDH | Rabbit | IB (1:5000) | sc-25778 | Santa Cruz<br>Biotechnology | FL-335 |
| Gbp2 | Goat | IB (1:200) | sc-10588 | Santa Cruz<br>Biotechnology | M15 |
| Gbp3 | Rabbit | IB (1:500) | This study |  | CLREEMERT<br>RRKPSLF |
| Gbp4 (mGbp3) | Goat | IB (1:200) | sc-242893 | Santa Cruz<br>Biotechnology | M17 |
| Gbp5 | Goat | IB (1:200) | sc-160335 | Santa Cruz<br>Biotechnology | G12 |
| Gbp7 | Rabbit | IB (1:1000) | This study |  | CQMNQTQN<br>SDKVRKL |
| GFP | Rabbit | IB (1:5000) | A01704 | Genscript |  |
| Goat anti-Rat IgG (H+L)-<br>HRP | Rat | IB (1:1000) | 31470 | Thermo Fisher |  |
| Gsdmd | Rabbit | IB (1:1000) | ab209845 | Abcam | EPR19828 |
| HA.11 | Mouse | IB (1:10000),<br>IF (1:1000) | 901501 | BioLegend |  |
| IgG (Goat)-Alexa488 or<br>594 or 647 | Donkey | IF (1:1000) | A11055 | Thermo Fisher | Anti-mouse |
| IgG (Mouse)-Alexa488 or<br>594 or 647 | Donkey | IF (1:1000) | A21203 | Thermo Fisher | Anti-goat |
| IgG (Rabbit)-Alexa488 or<br>594 or 647 | Donkey | IF (1:1000) | A21207 | Thermo Fisher | Anti-rabbit |
| IL-1β/IL-1F2 | Goat | IB (1:800) | AF-401 | R&D Systems | |
| Na<sup>+</sup>/K<sup>+</sup> ATPase | Mouse | IB (1:1000) | ab76020 | Abcam | EP1845Y |
| NLRP3/NALP3 | Mouse | IB (1:1000) | AG-20B-0006-C100 | Adipogen | Cryo-1 |
| Mouse IgG, HRP-linked<br>whole Ab | Mouse | IB (1:1000) | NXA931-1ML | GE Healthcare |  |
| Rabbit IgG, HRP-linked<br>whole Ab (from donkey) | Rabbit | IB (1:1000) | NA934-1ML | GE Healthcare |  |
| Streptavidin, Alexa Fluor®<br>647 Conjugate |  | IF (1:1000) | S32357 | Thermo Fisher |  |

